# Focusing antibody responses to the fusion peptide in rhesus macaques

**DOI:** 10.1101/2023.06.26.545779

**Authors:** Christopher A. Cottrell, Payal P. Pratap, Kimberly M. Cirelli, Diane G. Carnathan, Chiamaka A Enemuo, Aleksandar Antanasijevic, Gabriel Ozorowski, Leigh M. Sewall, Hongmei Gao, Kelli M. Greene, Joel D. Allen, Julia T. Ngo, Yury Choe, Bartek Nogal, Murillo Silva, Jinal Bhiman, Matthias Pauthner, Darrell J. Irvine, David Montefiori, Max Crispin, Dennis R. Burton, Guido Silvestri, Shane Crotty, Andrew B. Ward

**Affiliations:** Department of Integrative Structural and Computational Biology, The Scripps Research Institute, La Jolla, CA 92037, USA; International AIDS Vaccine Initiative Neutralizing Antibody Center, the Collaboration for AIDS Vaccine Discovery (CAVD) and Scripps Consortium for HIV/AIDS Vaccine Development (CHAVD), The Scripps Research Institute, La Jolla, CA 92037, USA; Center for Infectious Disease and Vaccine Research, La Jolla Institute for Immunology, La Jolla, CA 92037, USA.; Division of Microbiology and Immunology, Emory National Primate Research Center, Emory University, Atlanta, GA 30329, USA; Duke Human Vaccine Institute and Department of Surgery, Duke University Medical Center, Durham, NC, USA; School of Biological Sciences, University of Southampton, Southampton, SO17 1BJ, UK; Department of Immunology and Microbiology, The Scripps Research Institute, La Jolla, California, USA; Koch Institute for Integrative Cancer Research, Massachusetts Institute of Technology, Cambridge, MA 02139, USA; Centre for HIV and STI, National Institute for Communicable Diseases of the National Health Laboratory Service, Johannesburg, South Africa; Ragon Institute of MGH, MIT and Harvard, Cambridge, MA02139, USA; Division of Infectious Disease and Global Public Health, Department of Medicine, University of California, San Diego, La Jolla, California, USA

## Abstract

Immunodominance of antibodies targeting non-neutralizing epitopes and the high level of somatic hypermutation within germinal centers (GCs) required for most HIV broadly neutralizing antibodies (bnAbs) are major impediments to the development of an effective HIV vaccine. Rational protein vaccine design and non-conventional immunization strategies are potential avenues to overcome these hurdles. Here, we report using implantable osmotic pumps to continuously deliver a series of epitope-targeted immunogens to rhesus macaques over the course of six months to elicit immune responses against the conserved fusion peptide. Antibody specificities and GC responses were tracked longitudinally using electron microscopy polyclonal epitope mapping (EMPEM) and lymph node fine-needle aspirates, respectively. Application of cryoEMPEM delineated key residues for on-target and off-target responses that can drive the next round of structure-based vaccine design.

## Introduction

Most licensed vaccines are delivered via an initial bolus priming immunization followed by subsequent booster immunizations with the same antigen^1^. These booster immunizations are given to increase the titers of neutralizing antibodies and provide an additional opportunity for protection in individuals that did not respond to the priming immunization^1–4^. Conversely, all HIV broadly neutralizing antibodies (bnAbs), which are endowed with the most desirable properties for protection against diverse HIV strains and have demonstrated prophylaxis against infection in recent human trials, develop in response to constant exposure to a continuously evolving antigen, envelope (Env), in the context of long-term infection^5–7^. This continuous stimulation of germinal centers (GCs) drives high amounts of somatic hypermutation required for bnAb evolution^8–14^. We hypothesized that the immune system may therefore be optimized to generate bnAbs in GC responses in the presence of continuous antigen, as is the case during chronic infection, rather than episodic antigen exposure (e.g. conventional bolus needle immunization). Indeed, short duration (1 to 2 weeks) antigen delivery using subcutaneously (s.c.) implanted osmotic pumps has been used in mice and rhesus macaques (RMs) to protect immunogens from *in vivo* degradation and facilitate slow release of antigen to stimulate GCs for enhanced germinal center B (B_GC_) cell and T follicular helper (Tfh) cell responses^15–18^. In all cases, animals immunized via osmotic pump had better overall immune responses compared to control animals receiving bolus immunizations^15–18^.

Multiple sites of vulnerability have been identified on the Env glycoprotein against isolated bnAbs, including the CD4 binding site (CD4bs), the V2-apex, the V3-glycan site, the membrane proximal external region (MPER), the gp120/gp41 interface and the fusion peptide (FP)^9,11–13,19–22^. The highly conserved FP is necessary for viral fusion into host cells and has been shown to be a broadly reactive site of vulnerability on Env in HIV vaccine efforts^23–25^. Previous work in RMs has shown that recombinant BG505 SOSIP Env trimers are capable of eliciting autologous, and in some cases weakly heterologous, neutralizing antibodies directed to the FP epitope^26–28^. Prime-boost regimes employing FP scaffold immunogens and the BG505 SOSIP trimer also have elicited FP-directed neutralizing antibodies in multiple animals, including some with neutralization breadth^29,30^. While this approach shows promise, inducing consistent responses with fewer immunizations will be necessary for an effective vaccine.

Here, we combined a BG505-based immunogen series engineered to prime and mature antibody responses to the FP epitope with a platform for continuous immunogen delivery over the course of six months in non-human primates (NHPs). Further, we sought to recapitulate some aspects of the immune response during chronic infections by varying the immunogen over the course of the continuous delivery. Using electron microscopy polyclonal epitope mapping (EMPEM), we carefully monitored the antibody responses over time to evaluate our strategy.

## Results

### Design of a fusion peptide epitope targeting immunogen

The stabilized BG505 SOSIP.v5.2 was used as the basis for engineering a series of trimer immunogens designed to focus antibody response to the FP epitope^31^. For the priming immunogen, we removed the N611 glycan to make the FP epitope more accessible to B cell receptors (BCRs) and added a potential N-linked glycosylation site (PNGS) at N289 to reduce off-target antibody responses directed to the N289 glycan hole (Fig. 1A)^32–34^. In the Boost#1 immunogen we added a PNGS at N241 to restrict the angles of approach for BCRs to target the FP epitope in a bnAb-like fashion (Fig. 1A). Subsequently, the Boost#2 immunogen was designed to restore the N611 PNGS with an additional S613T mutation that enhances glycan occupancy at the N611 site (Fig. 1A)^35^. The BG505 Env sequence has several residues surrounding the FP epitope that are poorly conserved when compared to the Los Alamos National Library (LANL) HIV Sequence database (https://www.hiv.lanl.gov/content/sequence/HIV/HIVTools.html); particularly, H85 (8.6% prevalence among global strains) and K229 (13.0% prevalence), which have been shown to play a role in the epitopes of autologous nAbs elicited by BG505 SOSIP in rabbits and RMs^26,36^. Hence, for the Boost#3 immunogen, we mutated H85 and K229 to the consensus residues valine (39.0% prevalence) and asparagine (78.6% prevalence), respectively (Fig. 1A). The antigenicity of all four immunogens was evaluated by biolayer interferometry using a panel of anti-HIV Env antibodies (Figs 1A and S1). All four immunogens bound well to the quaternary specific bnAb PGT145 and showed little binding to the gp120 (monomer)-specific F105 antibody (Fig. 1B)^37–39^. All four immunogens showed low binding to the V3 targeting non-nAbs 19b and 14e, no detectable binding to a panel of non-nAbs including those targeting the gp41 post-fusion conformation (F240, 7B2, and 98-6), and high binding to the BG505-SOSIP elicited base-binding mAb RM19R (Fig. S1A)^26^. Additionally, all four immunogens bound to known FP targeting bnAbs (PGT151, VRC34, and ACS202) (Fig 1B)^23,25,40^. The Prime immunogen bound to an inferred germline (iGL) version of FP-targeting RM20F, originally isolated from an NHP immunized with BG505 SOSIP (Fig 1B)^26^. The iGL-RM20F mAb bound weakly to the Boost#1immunogen and did not bind to the Boost#2 or Boost#3 immunogens (Fig 1B). The mature version of RM20F, which can potently neutralize autologous virus, also exhibited diminished binding to the later boost immunogens (Fig 1B)^26^. Hence, the Prime and Boost#1 immunogens were designed to engage a known FP BCR while the Boost#2 and #3 were designed to mature toward breadth and away from the strain specific phenotype embodied by RM20F.

**Figure 1.**
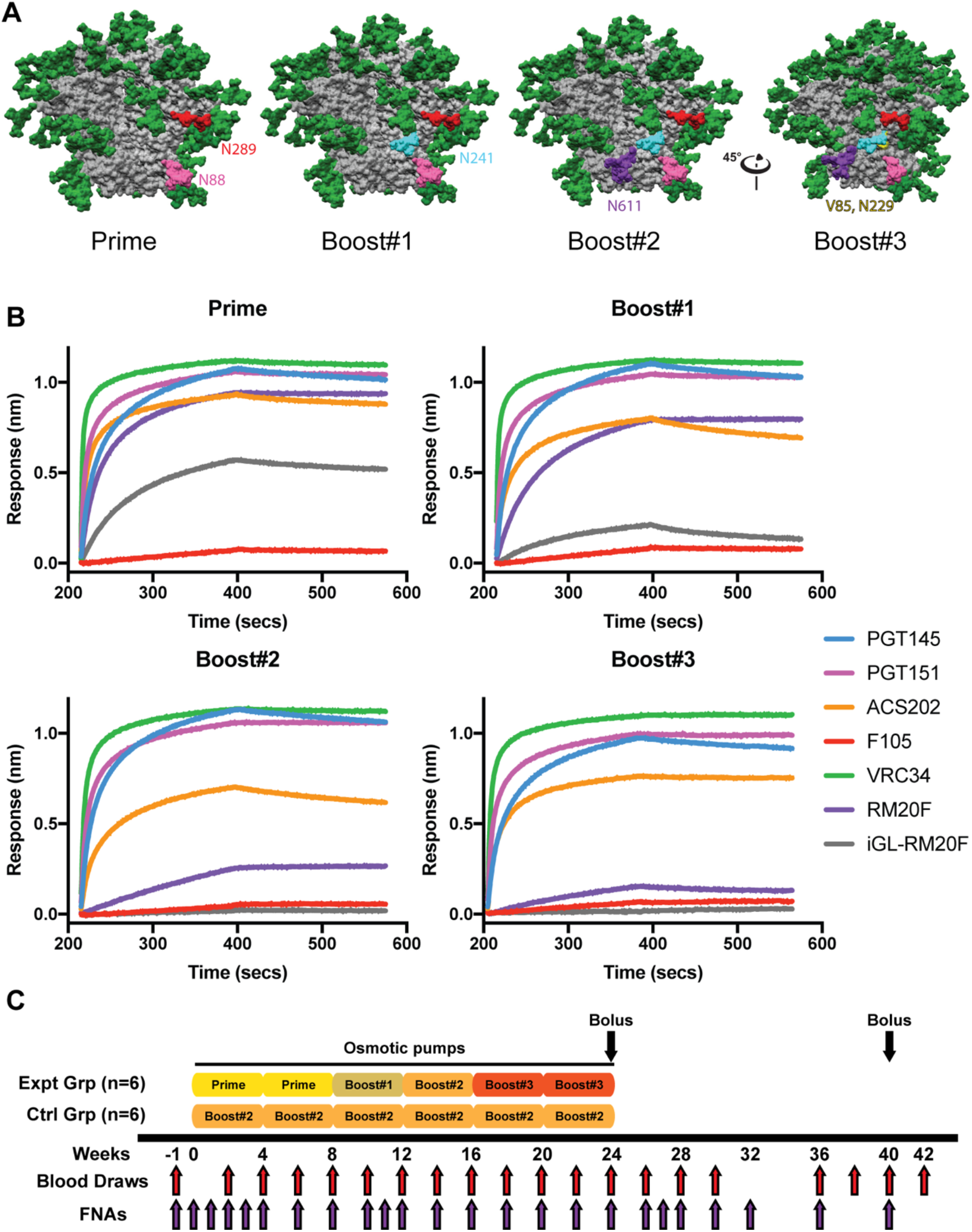
FP focused immunogen design and immunization scheme. **(A)** Surface representations of the BG505 SOSIP.v5.2 immunogen with mutations designed to focus antibody responses to the FP epitope. N88 glycan in pink. N611 glycan in purple. Introduced N289 glycan in red. Introduced N241 glycan in turquoise. Consensus mutations around the FP epitope shown in yellow. **(B)** Antigenicity was assessed using BLI on panel of anti-Env mAbs: PGT145 (apex), PGT151 (FP), ACS202 (FP), VRC34 (FP), RM20F (FP), iGL-RM20F (FP), and F105 (anti-gp120, non-Nab). **(C)** Immunization scheme.

Two groups of six RMs each were immunized continuously over the course of 24 weeks using subcutaneously (s.c.) implanted osmotic pumps (Fig. 1C). The pumps were filled with immunogen adjuvanted with Matrix-M (which has been successfully incorporated in the approved Novavax protein subunit COVID vaccine^41^) and implanted bilaterally in the left and right thighs, with exchanges occurring every 4 weeks^42^. The experimental group received the Prime immunogen in the first two pumps, the Boost#1 immunogen in the 3^rd^ pump, the Boost#2 immunogen in the 4^th^ pumps and Boost#3 in the final two pumps (Fig 1C). The control group received the Boost#2 immunogen in all six pumps (Fig. 1C). At the conclusion of the continuous immunogen delivery portion of the study, the animals were given an s.c. bolus immunization with an octameric nanoparticle immunogen displaying 8 copies of either the Boost#3 or Boost#2 immunogens for the experimental and group controls, respectively. The nanoparticle immunization was given bilaterally in the left and right thighs with the 3M-052 stable emulsion (SE) adjuvant^43^. The animals were boosted again at week 40 via s.c. bolus immunization using SMNP adjuvant with either the Boost#3 or Boost#2 soluble trimer immunogens for the experimental and control groups, respectively (Fig 1C)^44^.

### GC responses were durable over the course of six months continuous immunization

To monitor the GC responses during continuous immunogen delivery, longitudinal lymph (LN) node fine needle aspirates (FNAs) were used to sample the draining inguinal LNs throughout the study (Fig 1C). Previous work has shown that LN FNAs are well tolerated and represent the cellular composition of the whole LN^15,45^. B_GC_ cell frequencies rose during the course of the first pump and remained elevated throughout the study (Fig. 2A-B). Cells taken from the LN FNAs were stained with fluorescent probes of the Prime and Boost#2 immunogens. Overall, the frequency of B_GC_ cells that were double positive for both immunogens was consistent between groups throughout the study (Fig. 2C-F). The frequency of B_GC_ cells that were positive for only the Prime immunogen was significantly greater in the experimental group, consistent with a bias toward the gp41-N611/FP region. (Fig. 2E-H).

**Figure 2.**
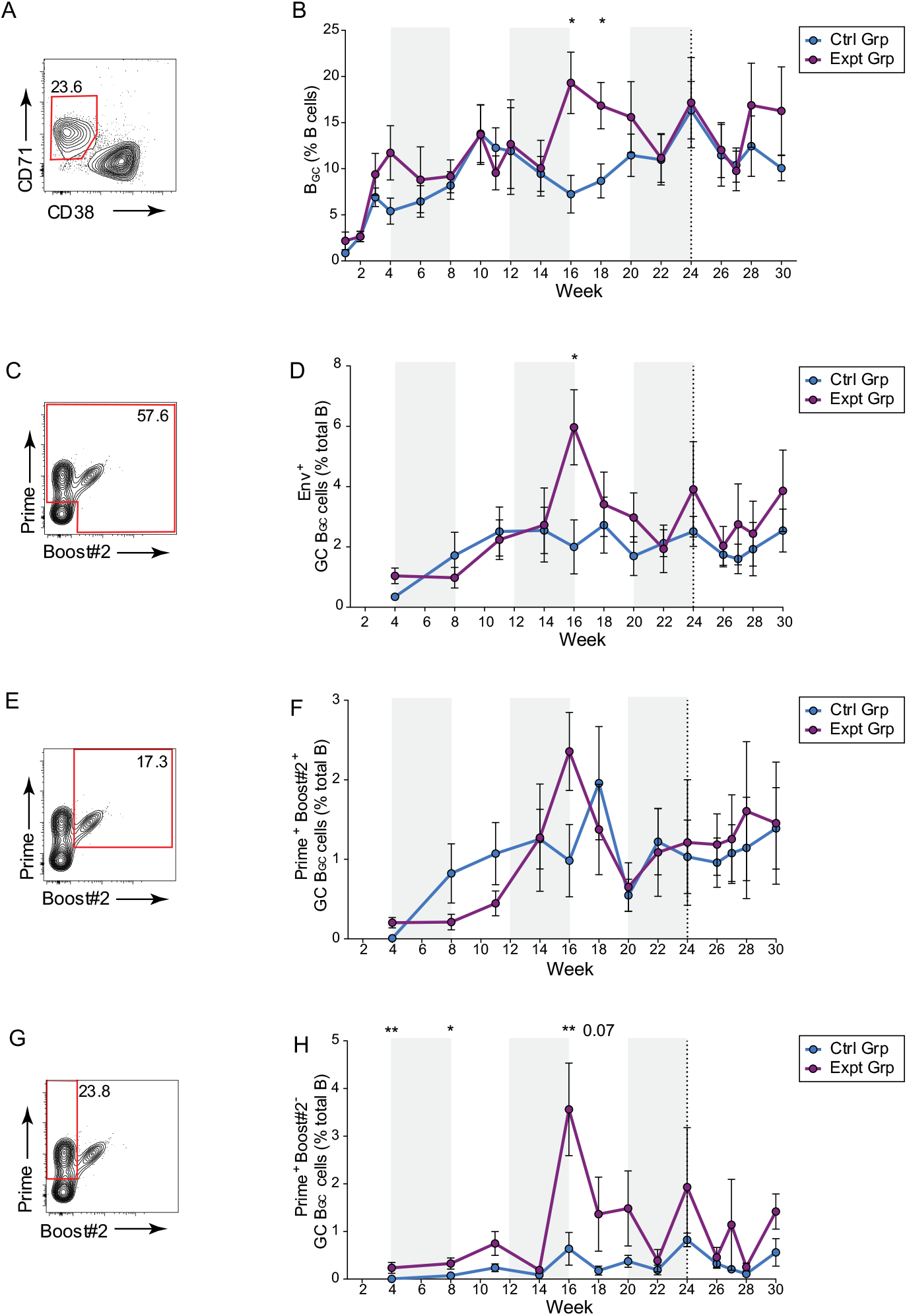
Sustained delivery immunization results in enduring GC responses. **(A)** Representative B_GC_ cell flow cytometry. **(B)** Quantification of B_GC_ cell kinetics as a percentage of total CD20^+^ B cells. **(C)** Representative total Env trimer-specific binding flow cytometry, gated on B_GC_ cells. **(D)** Quantification of total Env trimer-specific B_GC_ cells kinetics as a percentage of total CD20^+^ B cells. (**E)** Representative Prime and Boost#2 dual-specific binding flow cytometry, gated on B_GC_ cells. (**F**) Quantification of Prime and Boost#2-specific cell kinetics as a percentage of total CD20^+^ B cells. **(G)** Representative Prime immunogen-specific flow cytometry, gated on B_GC_ cells. **(H)** Quantification of Prime immunogen-specific B_GC_ cell kinetics as a percentage of total CD20^+^ B cells. Plot shows the mean of left and right side FNAs (analyzed separately) with standard error of means (SEM), * p ≤0.05, **p ≤0.01.

### Longitudinal monitoring of humoral immune responses revealed early targeting of the FP epitope in the experimental group

To assess antibody specificities elicited by immunization, polyclonal IgG isolated from plasma blood draws was digested into fragments antigen binding (Fabs) and subjected to EMPEM analysis^46^. Prior to week 18, EMPEM analysis was conducted using either the Prime (expt group) or Boost #2 (ctrl group) immunogens ad probes, and the results are reported as composites of both runs. Polyclonal Fabs isolated as early as week 6 from the experimental group targeted the gp41-N611/FP region (Fig. 3). In both immunization groups, antibodies directed to the base of the trimer were observed as early as week 6 and persisted throughout the duration of the study (Figs. 3C and 3D), consistent with immunodominance of the base epitope^16,26,47,48^. Antibody responses directed to non-FP, non-neutralizing epitopes other than the base of the trimer (V5/C3, V1/V3, gp120/gp120 interface (IF), and N355/N289 glycan hole epitope) were observed earlier and more consistently in the control group during the continuous immunogen delivery phase of the study (Fig. 3C). By week 14 (2 weeks into the pump containing the Boost#2 immunogen), antibody responses directed towards these off-target epitopes became prevalent in the experimental group (Fig. 1C). This delay in eliciting antibodies directed to off-target epitopes other than the base of the trimer in the experimental group suggests the gp41-N611/FP region may have been preferentially targeted relative to the others. By the end of the continuous immunogen delivery phase both groups of animals had a similar prevalence of antibodies directed to off-target epitopes with the experimental group also maintaining a high prevalence of gp41-N611/FP region directed antibodies (Fig. 3C).

**Figure 3.**
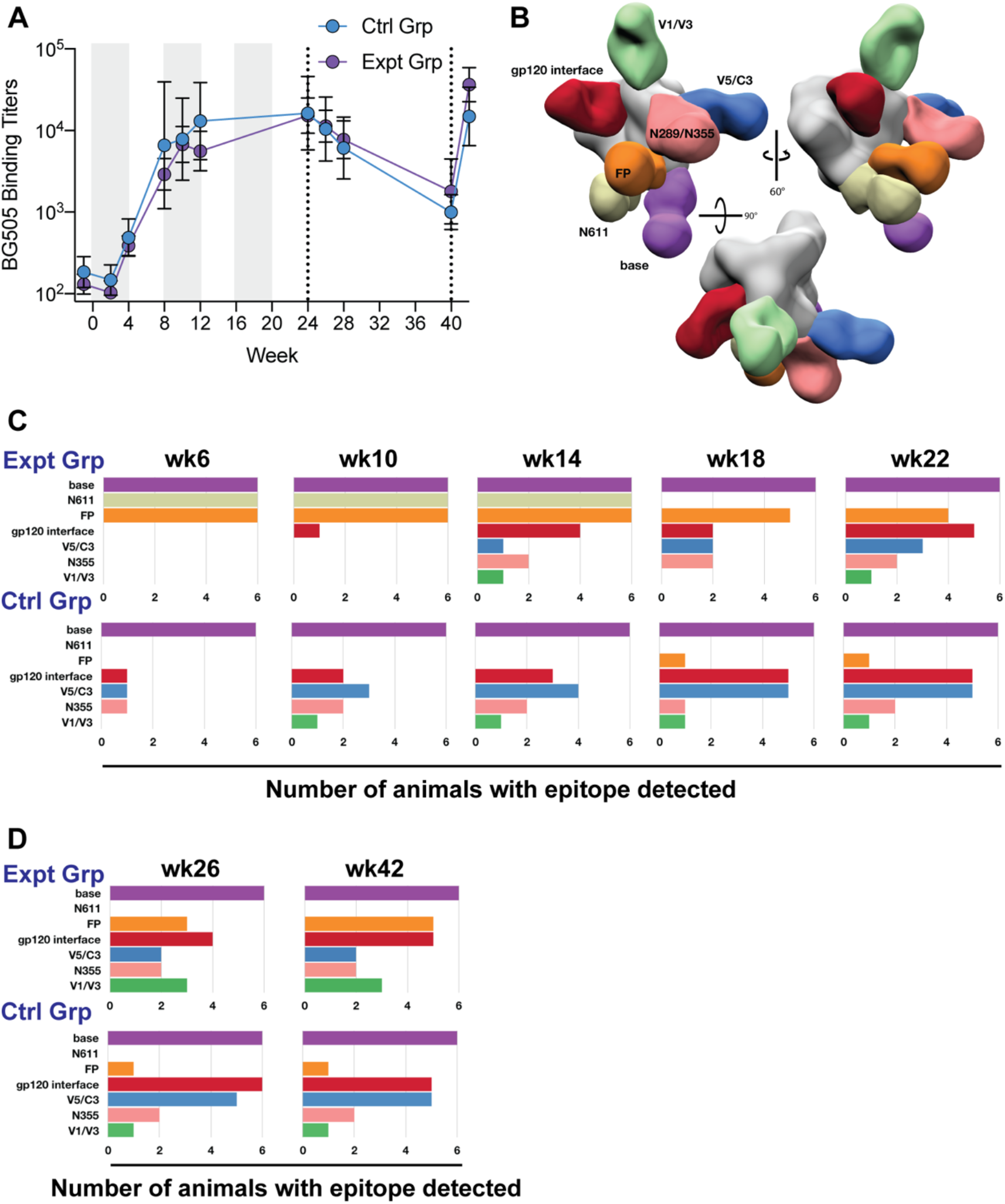
Longitudinal monitoring of the humoral immune response by ELISA and negative stain EMPEM. **(A)** Plasma IgG binding ELISA of each group over the duration of the study. Six animals per group, showing geometric mean titers ± geometric SD. **(B)** Composite 3D map representing the epitopes observed in the longitudinal negative stain EMPEM analysis. **(C)** Longitudinal negative stain EMPEM analysis of the continuous immunogen delivery phase of the study. Bar graphs show how many animals per group made antibodies directed to each specific epitope. Colored to match (**B**). **(D)** Longitudinal negative stain EMPEM analysis 2 weeks after each bolus immunization, colored to match (**B**).

The bolus immunizations with the octameric immunogen did not boost the BG505 Env binding titers in either group (Fig. 3A) nor did they have a detectable impact on the epitopes visualized by EMPEM analysis (Fig. 3D). Instead, the octameric immunogen appeared to have induced a de novo antibody response directed to the protein core of the nanoparticle (Fig. S2). Compared to the end of the continuous pump and octameric immunogen bolus immunizations, week 40 bolus immunizations with trimer immunogen and SMNP led to a bigger increase of BG505 binding titers in the experimental group compared to the control groups (Fig. 3A) with no detectable impact on the epitope diversity seen in the EMPEM analysis in either group (Fig. 3D).

At the end of the continuous immunogen delivery phase, the BG505 autologous neutralization titers were lower in the experimental groups compared to the control group (Fig. 4A). The bolus immunization with the octameric immunogen did not improve the BG505 autologous neutralization titer for either group (Fig. 4B, Table S2). Sera from animals in the experimental group potently neutralized the BG505 pseudovirus with the N611 glycan knocked out (N611A) (Fig. 4C, Table S2). This enhancement in neutralization activity following the removal of the N611 glycan was absent in the control group (Fig. 4C, Table S2). Adding a glycan to the V5 loop (T465N) decreased the neutralization activity of the sera from the control group (Fig. 4D, Table S2), consistent with the V5 epitope seen in the EMPEM analysis which was likely responsible for the autologous neutralization activity. Additionally, lengthening the V1 by two amino acids (133aN and 136aA), which has been previously shown to be part of a neutralization epitope targeted in BG505 SOSIP animals, decreased the neutralization activity of sera from animals 31943 (Ctrl Grp) and 32607 (Expt Grp), both of which had V1 epitope targeting antibodies detected by EMPEM analysis (Figs 3B-3D)^27,49,50^. Neither group demonstrated significant neutralization breadth when assessed on the global panel or an FP-sensitive panel of pseudoviruses (Tables S2 – S3)^30,51^. BG505 neutralization titers significantly increased after trimer and SMNP bolus immunizations (week 42) compared to after octameric bolus immunizations (week 26) (Fig. 4F) (Table S2). Week 42 FP-sensitive neutralization titers showed two of the animals in the experimental group were able to weakly neutralize three out of nine pseudoviruses in the FP sensitive panel while none of the animals in the control group were able to neutralize any of the pseudoviruses in the FP-sensitive panel at this time point (Table S3). Further, in-depth epitope mapping analysis was done on the animal with higher neutralization titers, which is discussed later.

**Figure 4.**
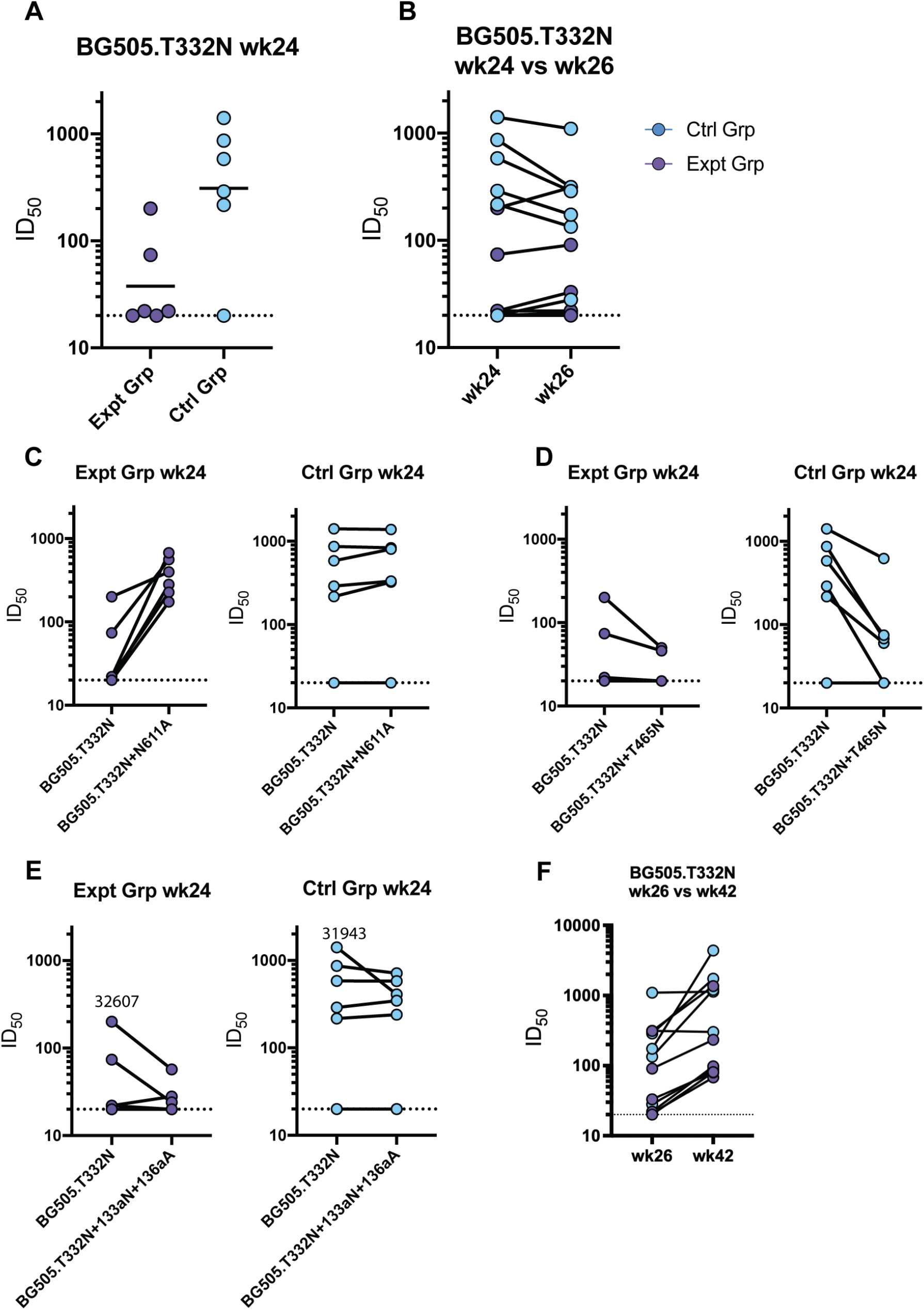
Serum neutralization activity. **(A)** Week 24 serum neutralization activity against the BG505.T332N pseudovirus. Dotted lines indicate detection limit for the assay. **(B)** Comparison of BG505.T332N pseudovirus neutralization activity against before (week 24) and after (week 26) bolus nanoparticle immunization. Connecting lines indicate specific animal response differences seen between the two time points. **(C)** Week 24 serum neutralization activity against the BG505.T332N+N611A pseudovirus. **(D)** Week 24 serum neutralization activity against the BG505.T332N+T465N pseudovirus. **(E)** Week 24 serum neutralization activity against the BG505.T332N+133aN_136aA pseudovirus. **(F)** Week 26 versus week 42 serum neutralization activity against BG505.T332N pseudovirus.

At the week 18 time point, we transitioned to using only the Boost#2 immunogen as a probe for both groups. This was done to detect FP directed antibodies that could accommodate the N611 and N241 glycans present in the Boost#2 immunogen. As expected, the use of this trimer resulted in the inability to detect N611 glycan hole-directed antibody responses in the experimental group after week 18 (Fig. 3C and 3D).

Surprisingly, several animals in both groups made antibody responses directed to the N355/N289 glycan hole epitope, despite the introduction of the N289 PNGS sequon in all the immunogens used (Fig. 3C and 3D). Post-hoc, site-specific glycan occupancy analysis was conducted on the immunogens (Fig. S3 to S7). Unfortunately, the analysis could not detect peptides corresponding to the N289 site and glycan occupancy at that site could not be measured. Given the heterogenous occupancy of the introduced N241 PNGS in the Boost#1, Boost#2 and Boost#3 immunogens, along with the EMPEM data, the N289 glycan occupancy was likely less than 100% in the immunogens and flow cytometry probes.

### CryoEMPEM elucidation of epitope/paratope interfaces of polyclonal Fabs elicited through immunization against the Boost#2 Immunogen

#### Fusion Peptide

CryoEMPEM (EMPEM using vitrified instead of negatively-stained samples) can resolve different classes of antibodies with overlapping binding sites within one sample and provide a more detailed picture of epitope/paratope interfaces compared to negative stain (ns) EMPEM^47^. The nsEM analysis of animal Rh.32613-week 42 protein complex revealed interesting FP-directed responses and was chosen for cryoEMPEM analysis. 13 distinct, high resolution (≤4.5 Å) antibody-immunogen maps were reconstructed from this single dataset, including four different antibodies against the FP region (FP1-4) (Fig. S5 and Table S1). Of the four maps, FP1 and FP3 had the most interpretable density such that atomic modeling could be attempted.

The FP3 map (∼3.5 Å resolution) contains a fully resolved N-terminus of the FP, which due to the inherent flexibility of the hydrophobic linear peptide, is typically resolved only via cryoEM when stabilized by an anti-FP antibody as seen in FP3 and bnAbs isolated from humans (Figs. 5A and 5B)^23,25,30,52–54^. In the FP3 complex conformation, the FP N-terminus is stabilized via interactions with L645 (90.2% prevalence) and E648 (58.6% prevalence) of the adjacent gp41 subunit and residue 69 of the light chain framework region 3 (LFR3) (Fig 5C and 5D). Atomic modeling of the FP3 antibody backbone revealed a long heavy chain complementarity determining region 3 (HCDR3) with a predicted length of 25 amino acids (aa), which is a common feature of bnAbs (Fig S5A and Table 1)^13,55^. The densities of the HCDR3 side chains suggest the presence of at least three aromatic residues and other medium-sized hydrophobic residues interacting with the fusion peptide (Fig. 5D) ^56^. A hydrophobic pocket forms around a predicted aromatic residue at position 101 of the heavy chain, presumed to be a tryptophan based on the density, which interacts with the I515 (53.0% prevalence) and V518 (43.2% prevalence) residues of the FP as well as I641 (19.4% prevalence) on the heptad repeat domain 2 (HR2) of gp41. Another hydrophobic pocket is found around F522 of the FP, which is further stabilized by the presence of two aromatic residues at positions 107 and 108 of the HCDR3 (Fig. 5D). The long HCDR3 also interacts with many residues of the C1 region of gp120, including highly conserved residues T77 (98% prevalence), D78 (98.7% prevalence), Q82 (prevalence 94.0%), E83 (99.5% prevalence) as well as less conserved residues I84 (45.2%), H85 (8.6%) and E87 (56.1%).

**Figure 5.**
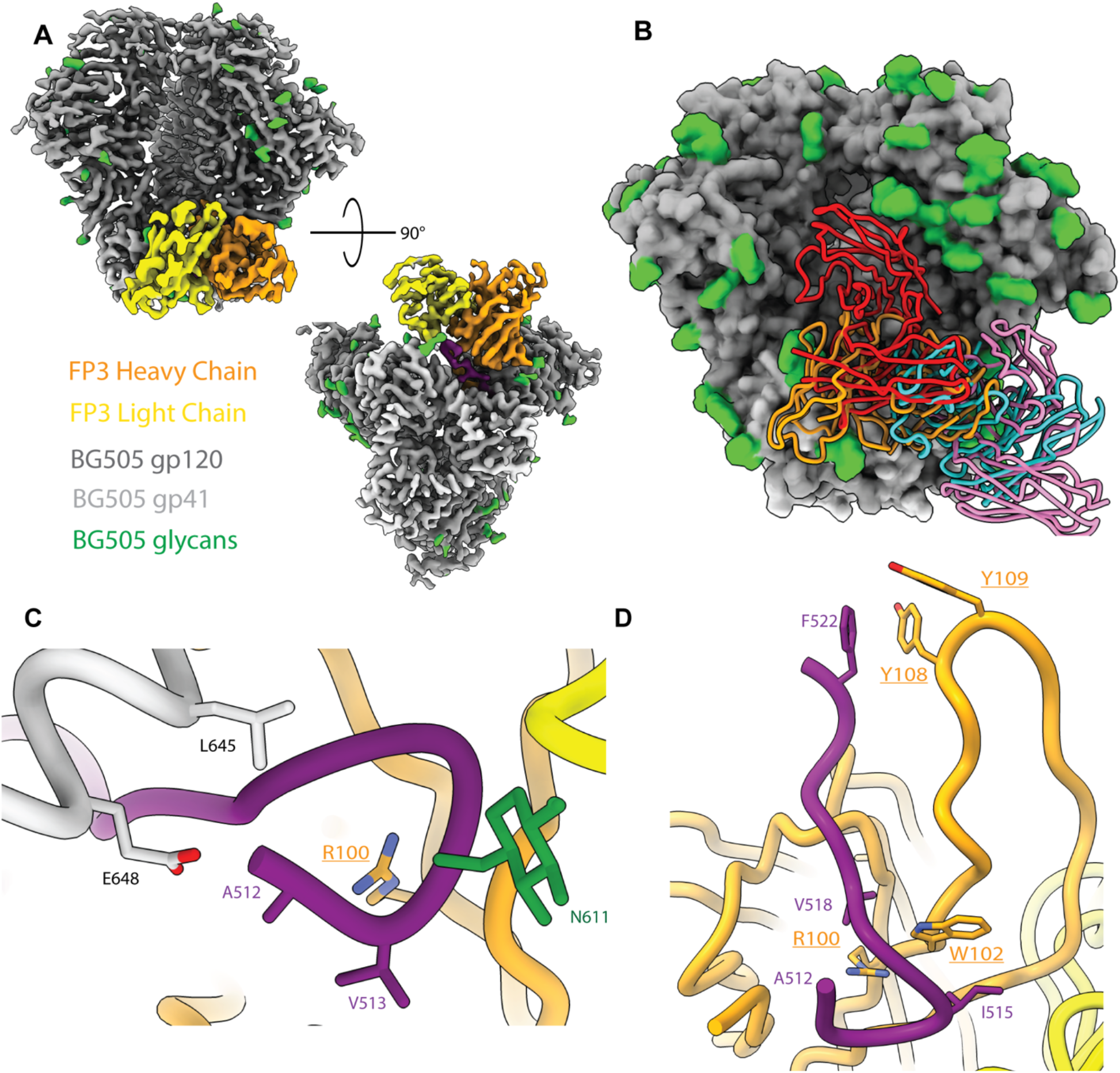
CryoEMPEM Analysis of FP3. A) Boost #2 in complex with the predicted heavy chain (orange) and light chain (yellow) of FP3 antibody targets the fusion peptide epitope area. B) Alignment of FP3 (orange) compared to known human bnAbs PGT151 (red), VRC34.01 (cyan), and ACS202 (pink). C. Fusion peptide is fully resolved in FP3 due to stabilization of the N-terminus by the antibody (underlined residues are inferred from density; residues not underlined are sequence verified). D) The CDRH3 of FP3 antibody interacts with FP by creating hydrophobic pockets around the N-terminus as well as around F522 with the presence of hydrophobic aromatic residues.

**Table 1.** Fusion peptide nAbs. Summary of isolated human or NHP nAbs against the fusion peptide, including the FP contacts, non-FP contacts, glycan contacts and, VH gene usage and HCDR3 lengths. Contacts are defined as a distance <5 Å between antibody and HIV Env trimer atoms.

| FP Abs<br>(PDB Code) | Species | Antigen<br>Genotype | FP aa<br>contacts | Non-FP<br>aa<br>contacts | Glycan<br>contacts | VH Gene | HCDR3<br>length<br>(aa) |
| --- | --- | --- | --- | --- | --- | --- | --- |
| VRC34.01<br>(5I8H) | Human | BG505 | 512-519 | I84, H85,<br>K229 | N88 | IGHV1-2*02 | 13 |
| (6NC3) |  | AMC011 | 512-520 | V85 | N88,<br>N241 |  |  |
| PGT151<br>(6OLP) | Human | AMC011 | 512-522 | P79,<br>Q82,<br>E83,<br>V84,<br>Q246,<br>R542-L543<br>S640,<br>Y643,<br>T644 | N611,<br>N637 | IGHV3-30*04 | 28 |
| ACS202<br>(6NC2) | Human | AMC011<br>v4.2 | 512-520,<br>A525 | V85,<br>E87,<br>S528,<br>A532, | N88,<br>N241 | IGHV3-30*03 | 24 |
|  |  |  |  | A533,<br>M535,<br>T536 |  |  |  |
| RM20F<br>(6VN0) | Macaque | BG505 | 518,<br>520-521 | P81, E83,<br>I84, H85,<br>E87,<br>K229,<br>K231,<br>M535,<br>T536,<br>V539,<br>R542,<br>Q650 | N88 | IGHV3-AFY*10 | 20 |
| FP1 | Macaque | BG505 | 517-518,<br>520-521, 524 | D78,<br>N80-H85,<br>E87,<br>K229,<br>A532,<br>M535-T536,<br>T538-V539,<br>R542,<br>Q640, | N88,<br>N241 | LJI.Rh_IGHV3<br>.76.a_S4190 | 18* |
|  |  |  |  | G644,<br>E647-E648 |  |  |  |
| FP3 | Macaque | BG505 | 512-522 | P76-P79,<br>E82-H85,<br>E87,<br>A219-I225,<br>T244-Q246,<br>I491,<br>R542-L544,<br>S546,<br>T613-W614,<br>Y638,<br>Q640-I641,<br>Y643 | N88,<br>N637,<br>N611 | IGHV4-AFI*01 | 25* |
| DF1W-a.01<br>(6MPH) | Macaque | BG505<br>DS-SOSIP | 512-520 | N80,<br>P81,<br>Q82,<br>E83, I84,<br>E87, | N88 | IGHV4-AFQ*01_S086<br>8 | 10 |
|  |  |  |  | V539,<br>R542,<br>G644,<br>E648 |  |  |  |
| A12V163-a.01 (6N1V) | Macaque | BG505<br>DS-SOSIP | 512-519 | N80,<br>E83, I84,<br>H85,<br>E87, N88 | N88 | IGHV4-AFB*01 | 14 |
| A12V163-b.01 (6MPG) | Macaque | BG505<br>DS-SOSIP | 512-519 | D78,<br>N80,<br>Q82,<br>E83, I84,<br>H85,<br>E87,<br>K229,<br>A532,<br>R542,<br>Q640 | N88 | IGHV3-ADR*01 | 11 |
| DFPH-a.15<br>(6N1W) | Macaque | BG505<br>DS SOSIP | 512-520 | N80,<br>P81,<br>Q82,<br>E83, I84,<br>H85,<br>V539,<br>R542, |  | IGHV4-AGR*01_S409<br>9 | 19 |
|  |  |  |  | Q640,<br>G644,<br>E647,<br>E648 |  |  |  |
| 0PV-b.01<br>(6OT1) | Macaque | BG505<br>DS SOSIP | 512-520 | N80,<br>M535,<br>S613,<br>Q640<br>G644,<br>E648 | N637 | IGHV4-AGR*01_S409<br>9 | 14 |

There are 4 canonical glycans around the FP at positions N88, N230, N241 and N611, some of which have been shown to be important for FP bnAb binding and affinity such as glycans at N88 and N241 for VRC34 and ACS202, respectively^23,25,30,53^. Wild type BG505 does not contain either N230 or N241 glycan sites, although the N241 glycan site was knocked-in for the boosting immunogens in this study. In the case of FP3, interactions were observed between N88 glycan and residues 76 and 77 of HFR3. On the adjacent gp41 of the stabilized FP, HFR3 residue 68 is predicted to interact with the N611 glycan and the T613 residue. While not a canonical fusion peptide glycan, the glycan at N637 on the adjacent gp41 also interacts with LCDR2. Using the Ward Lab rhesus macaque germline database (RhGLDb)^26^, we predicted the V_H_ of FP3 to be IGHV4-AFI*01, with up to four amino acid differences between our model and the germline genes (∼96% sequence identity), possibly due to SHM.

FP1(∼3.7 Å resolution) resembles RM20F in its angle of approach, with only a partially resolved FP starting at position G516, engagement with the N88 glycan, and contacts with the poorly conserved residues H85 and K229 in the C1 and C2 regions of gp120, respectively^26^ (Fig 6). Atomic modeling predicts that FP1 HCDR3 is approximately 18 residues long, two less than the CDRH3 for RM20F (Fig. 6B) (Table 1). The predicted V_H_ of FP1 using the RhGLDb is LJI.Rh_IGHV3.76.a_S4190 with up to 5 aa differences between the modeled sequence and this V_H_. RM20F, on the other hand, is predicted to use IGHV3-AFY*10 as its V_H_, indicating only 85.6% sequence similarity by amino acid (88.1% nucleotide) between the two predicted V_H_.

**Figure 6.**
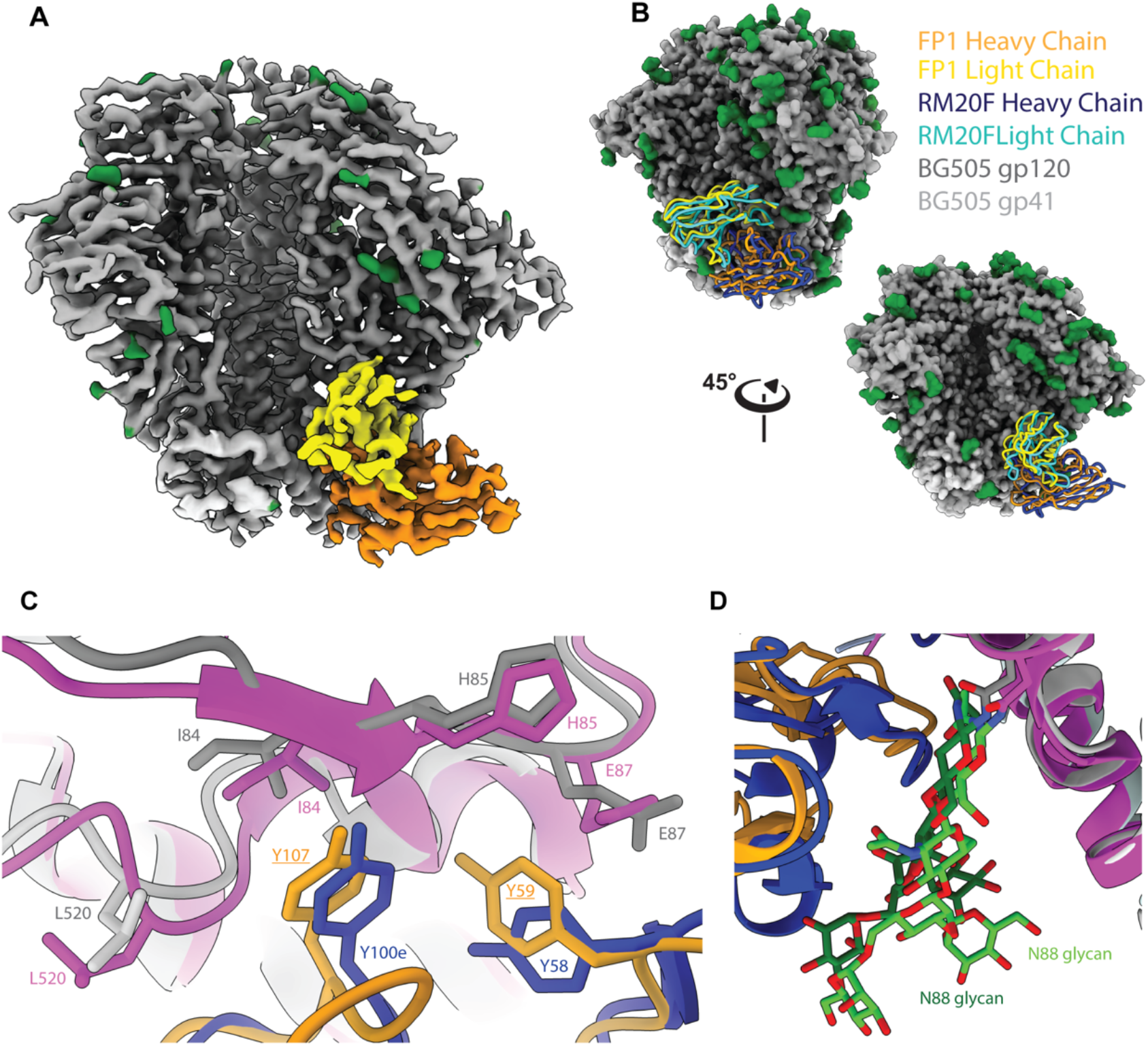
CryoEMPEM Analysis of FP1. A) Boost #2 in complex with predicted heavy chain (orange) and predicted light chain (yellow) of FP1. B) Alignment of FP1 (orange and yellow) with RM20F (dark and light blue) with two views. C) Both RM20F HCDR3 (dark blue) and FP1 predicted HCDR3 (orange) interact with I84 and H85 (grey for Boost #2 orientation for FP1 interactions; pink for BG505 residue orientation for RM20F (6VN0)) with a hydrophobic residue Y100e for RM20F and predicted Y106 (Y100c; Kabat) of FP1. CDRH2 of both FP1 and RM20F interact with H85 and E87 with hydrophobic residues at position 59 and 58, respectively. D) Heavy chains from both antibodies interact and stabilize the N88 glycan (dark green – FP1; neon green – 6VN0) for their respective antigen, allowing for branched resolution of the glycan.

#### Off-Target Responses

An additional four maps were also reconstructed for polyclonal responses against the gp41 base (Base1-4) (Fig. S4). Base-4 is the best resolved and demonstrates that residues R500 of gp120 and W623 of the intraprotomeric gp41 are important for this interaction. The predicted heavy chain for the pAb contains an HCDR3 that reaches into the base of the trimer and nestles between the R500 and W623 residues (Fig. S6A)^57,58^. Analysis of pAbs elicited against BG505 SOSIP base in other NHP studies also showed the prevalence of these residues within the base epitope/paratope interface^47^. The global prevalence of an arginine at position 500 on HIV Env is 24.8% but the prevalence of a positively charged residue (R, K, or H) at this position is 75.5%. The tryptophan clasp of gp41 (W623, W628 and W631) is critical for maintaining the prefusion confirmation of Env before protein rearrangement for viral entry upon engagement of the fusion peptide^57,58^. As a result, W623 is 99.5% conserved according to the LANL HIV Database. Two maps were derived that contained antibody responses to the gp120/gp120 IF epitope (Figs. S4), IF-1 and IF-3, each with sufficient resolution to explore the epitope/paratope interface. IF-1 Ab interacts with residues of the V3 loop including the stabilizing mutation A316W that was introduced beginning in SOSIP.v4 constructs to limit recognition by weakly/non-neutralizing antibodies (Fig. S6B)^31,59^. Antibody responses against this epitope have been reported in other NHP studies^47^. While the V3 loop and W316 residue are also in the epitope interface for IF-3 map, the antibody has additional interactions charged residues in the C1 and C2 regions including K65 (6.7% prevalence), H66 (99.9% prevalence) and K207 (99.3% prevalence) (Fig. S6C). Although we had introduced a glycan into position 289, which has a global prevalence of 70.0% for a PNGS at this position but is not present in wild type BG505, antibody responses were observed against theN289 residue, indicating the absence of the glycan at this position (Figs. S4 and S6C), opening this region for immune recognition.

## Discussion

Here we demonstrate that immunogen design was an effective means to successfully prime FP-directed antibody responses as well as the adjacent N611 glycan hole epitope. Over time however, and consistent with previous NHP slow delivery immunization studies, we observed a wide variety of epitopes targeted in this immunization protocol^15,28^. Despite introducing the N289 PNGS, typically missing in wild type BG505, into all the immunogens used in this study, several animals in both groups elicited antibodies against the N355/N289-glycan hole epitope. Glycan analysis was thus conducted on the immunogens to discern site specific glycan occupancy (Fig. S7 to S10). The analysis was not able to detect glycan occupancy at N289 site, but previous studies have shown that recombinant protein expression of Env trimer can have glycan under occupancy, even at highly conserved glycan sites^47^. For example, the introduced N241 PNGS in Boost#1, Boost#2 and Boost#3 immunogens showed differential levels of occupancy, so heterogenous occupancy for the N289 glycan would not be surprising and would explain the antibody evolution against this site. CryoEMPEM analysis of this epitope/paratope interface confirmed a lack of density corresponding to the N289 glycan, consistent with glycan under occupancy at this residue.

Presence of the V5/C3 epitope in EMPEM correlated with autologous neutralization activity that was subsequently reduced or eliminated by introducing a PNGS into the V5 loop at position 465^27,50^. The gp120/gp120-interface antibodies were not seen in previous EMPEM analysis in animals immunized with BG505 SOSIP.664, and based on previous results in RMs^26,30,47^, are thought to result from the A316W stabilizing mutation that falls within that epitope and is present in the BG505 SOSIPv5.2 construct used as the basis for immunogen design in this study.

CryoEMPEM further elucidated the FP and off-target responses. We were able to model two polyclonal antibodies, FP1 and FP3, that were able to either partially or fully stabilize the highly hydrophobic and flexible FP, respectively. FP3, with an estimated 25 aa HCDR3 not only fully engages with the FP but also makes contacts with other interprotomeric gp120 and gp41 regions. FP1, on the other hand, closely resembles RM20F, an antibody isolated from NHPs with autologous neutralization against BG505, in angle of approach, major contact residues and partial engagement with the FP.

Overall, this study shows that the FP/N611-glycan hole targeting antibodies can be reproducibly primed in RMs using an immune-focusing approach. GC response could be extended using continuous immunogen delivery but no gain in antibody neutralization breadth was observed relative to historical controls with short duration pumps or bolus immunizations. Hence, despite improved epitope specific responses and highly active and long-lived GCs, the NHPs failed to generate a broadly neutralizing antibody response. We therefore conclude that germline-targeting and additional immunogen design efforts are likely needed to improve the anti-FP antibody response and to elicit antibodies capable of neutralizing viruses with complete N611 glycosylation.

## Materials and Methods

### Rhesus Macaques

Indian origin rhesus macaques (*Macaca mulatta*) were sourced and housed at the Yerkes National Primate Research Center and maintained in accordance with NIH guidelines. This study was approved by the Emory University Institutional Animal Care and Use Committee (IACUC) [IACUC# 201700723]. When osmotic pumps were implanted, animals were kept in single, protected contact housing. At all other times, animals were kept in paired housing. Animals were treated with anesthesia and analgesics for procedures as per veterinarian recommendations and IACUC approved protocols. Animals were grouped to divide age, weight and gender as evenly as possible between groups.

### Immunogen and probe generation

The BG505 SOSIP.v5.2 and BG505 SOSIP.v5.2-Avitag plasmids were generously provided by Dr. Marit van Gils^31^. Site-direct mutagenesis (QuikChange Multi kit, Agilent) was used create the plasmids encoding the Prime, Boost#1, Boost#2, Boost#3, Prime-Avitag, and Boost#2-Avitag. Specific mutations relative to BG505 SOSIP.v5.2 are listed in Table S5. Immunogens and Avi-tagged probes were produced in HEK293F cells and purified using PGT145 affinity chromatography followed size-exclusion chromatography as described previously^59^. The Avi-tagged proteins were biotinylated using the BirA enzyme (Avidity) according to the manufacturer’s protocol. The resulting biotinylated proteins are referred to using the descriptor AviB.

### Immunizations

Osmotic pumps (Alzet model 2004) were loaded with 50 *μ*g soluble Env trimer immunogen + 75 U Matrix-M adjuvant (Novavax) in PBS according to Fig. 1C. Two pumps were subcutaneously implanted into each animal (one pump each in the left and right mid-thighs). The immunogen/adjuvant mixture was secreted continuously over the course of 4 weeks. The pumps were removed after 4 weeks and replaced with new pumps. At the conclusion of the sixth pump (week 24), the pumps were removed, and the animals received bilateral, bolus, s.c. immunizations with 59.6 *μ*g (118 *μ*g total dose) of an octameric nanoparticle^60,61^ formulated in 30 *μ*g 3M-052-squalene emulsion adjuvant (60 *μ*g total dose) in each mid-thigh. At week 40, the animals received additional bilateral, bolus, s.c. immunizations with 50 *μ*g (100 *μ*g total dose) soluble Env trimer immunogen formulated in 375 *μ*g (750 *μ*g total dose) SMNP adjuvant^44^ in each mid-thigh. Blood was collected at various time points into CPT tubes for PBMC and plasma isolation. Serum was isolated using serum collection tubes and frozen. Plasma was used in ELISA and EMPEM analysis. Serum was used for neutralization assays.

### Lymph node fine needle aspirates and FACS

Prime-AviB and Boost#2-AviB were individually premixed with fluorochrome-conjugated streptavidin (SA-Alexa Fluor 647 [Ax647] or SA-Brilliant Violet 421 [BV421]) at RT for 20 minutes. LN FNAs were used to sample at both right and left inguinal LNs. Draining lymph nodes were identified by palpitation and FNAs were performed by a veterinarian as described previously ^15,45^. Fresh cells from the FNAs were incubated with probes for 30 minutes at 4°C, washed twice with FACS buffer (PBS + 1% (v/v) Fetal bovine serum) and then incubated with surface antibodies (Table S6) for 30 minutes at 4°C. Cells were washed twice more with FACS buffer and sorted on an FACSAria II. B cells were defined as live CD20^+^CD4^-^CD8^-^CD16^-^, non-IgM^+^IgG^+^ cells. B_GC_ cells were further defined as CD71+/CD38− B cells.

### Monoclonal antibody production

Monoclonal IgGs were expressed in HEK293F cells and purified using affinity chromatography. Briefly, HEK293F cells (Invitrogen) were co-transfected with heavy and light chain plasmids (1:1 ratio) using PEImax. Transfections were performed according to the manufacturer’s protocol. Supernatants were harvested 4-6 days following transfection and passed through a 0.45 µm filter. IgGs were purified using MAbSelect™ (GE Healthcare) affinity chromatography.

### ELISAs

BG505 SOSIP IgG binding ELISA were conducted as described previously^62^. For the T33-31 ELISAs: Microlon half-area 96-well plates were coated overnight with 3 μg/mL T33-31 nanoparticle core protein in 0.1 M NaHCO_3_, pH 9. The plates were washed 3 times with wash buffer (TBS + 0.1% (v/v) Tween-20) and blocked overnight with PBS + 3% (w/v) BSA. The plates were washed 3 times with wash buffer and purified IgG from week 26 plasma was added in series for 5-fold dilutions starting at 1000 *μ*g/mL in PBS + 1% (w/v) BSA. The plates were incubated at RT for 2hrs and washed 3 times with wash buffer. Alkaline Phosphatase-conjugated AffiniPure Goat Anti-Human IgG/Fc secondary antibody (Jackson ImmunoResearch) was added at a 1:5000 dilution and allowed to bind for 1hr at RT. The plates were washed 4 times with wash buffer before being developed with 1-Step PNPP substrate (ThermoFisher) for ∼ 30 minutes. Absorbance at 405nm was recorded and data was analyzed with Prism version 8.4.2.

### TZM-bl cell-based neutralization assays – BG505 and FP-sensitive panels

Serum neutralization assays were conducted using the single-round infection assay of TZM-bl cells with HIV-1 Env-pseudoviruses as described previously^63–69^. Neutralizing antibodies were measured as a function of reductions in luciferase (Luc) reporter gene expression after a single round of infection in TZM-bl cells^64,65^. TZM-bl cells (also called JC57BL-13) were obtained from the NIH AIDS Research and Reference Reagent Program, as contributed by John Kappes and Xiaoyun Wu. This is a HeLa cell clone that was engineered to express CD4 and CCR5^66^ and to contain integrated reporter genes for firefly luciferase and E. coli beta-galactosidase under control of an HIV-1 LTR^67^. Briefly, a pre-titrated dose of virus was incubated with serial 3-fold dilutions of heat-inactivated (56°C, 30 minutes) serum samples in duplicate in a total volume of 150 µl for 1 hr at 37°C in 96-well flat-bottom culture plates. Freshly trypsinized cells (10,000 cells in 100 µl of growth medium containing 75 µg/ml DEAE dextran) were added to each well. One set of control wells received cells + virus (virus control) and another set received cells only (background control). After 48 hours of incubation, 100 µl of cells was transferred to a 96-well black solid plate (Costar) for measurements of luminescence using the Britelite Luminescence Reporter Gene Assay System (PerkinElmer Life Sciences). Neutralization titers are the dilution of serum samples at which relative luminescence units (RLU) were reduced by 50% compared to virus control wells after subtraction of background RLUs. Assay stocks of molecularly cloned Env-pseudotyped viruses were prepared by transfection in 293T/17 cells (American Type Culture Collection) and titrated in TZM-bl cells as described^64,65^. Mutations were introduced into Env plasmids by site-directed mutagenesis and confirmed by full-length Env sequencing by Sanger Sequencing, using Sequencher and SnapGene for sequence analyses. This assay has been formally optimized and validated^68^ and was performed in compliance with Good Clinical Laboratory Practices, including participation in a formal proficiency testing program^69^.

### TZM-bl cell-based neutralization assays – global panel

#### Envelope pseudovirus production

Envelope pseudoviruses were generated through the cotransfection of the pSG3ΔEnv backbone plasmid (obtained from the NIH AIDS Research and Reference Reagent Program, Division of AIDS, NIAID, NIH)^67,70^ and a plasmid encoding the full Env gp160 in a 3:1 ratio in HEK293T cells (ATCC) using the PEIMAX transfection reagent (Polysciences). Following 48 hours, the media was filtered through a 0.45 µm Steriflip unit (EMD Millipore), aliquoted, frozen and titrated. Mutations were introduced into Env plasmids by site-directed mutagenesis and confirmed by full-length Env sequencing by Sanger Sequencing, using Geneious Prime v8 for sequence analyses.

#### Neutralization assay

The neutralization assay^68^ was performed with DMEM (Corning) supplemented with 10% FBS (Corning), 1% L-glutamine (Corning), 0.5% Gentamicin (Sigma), 2.5% HEPES (Gibco) and all incubations were performed at 37 ° C, 5% CO2. On day one, 25 µL diluted pseudovirus mixed with 25 µL of 1:3 serially-diluted serum or control antibody was incubated for 1 hour, followed by the addition of 20 µL of TZM-bl cells at a concentration of 500, 000 cells/mL with DEAE-Dextran at a final concentration of 40µg/mL. These were incubated for 24 hours and on day two, 130 µL of fresh DMEM was added and the samples which were once again incubated overnight. Finally, on day three the media was completely removed and 60 µL lysis buffer together with 60 µL luciferase substrate (Promega) was added per well, and luminescence was measured on the Synergy2 plate reader (BioTek). Neutralization data is reported as ID50 or IC50 (µg/mL) values which was calculated as the dilution or concentration at which a 50% reduction in infection was observed. Neutralization assays were performed in triplicate and SEM is reported. The TZM-bl cell line engineered from CXCR4-positive HeLa cells to express CD4, CCR5, and a firefly luciferase reporter gene (under control of the HIV-1 LTR) was obtained from the NIH AIDS Research and Reference Reagent Program, Division of AIDS, NIAID, NIH (developed by Dr. John C. Kappes, and Dr. Xiaoyun Wu).

### Bio-Layer Interferometry (BLI)

An Octet RED instrument (FortéBio) was used to measure antibody–antigen interactions by Biolayer Interferometry. MAbs were loaded onto anti-human Fc (AHC) biosensors (FortéBio) at a concentration of 5 μg/mL in kinetics buffer (PBS, pH 7.4, 0.01% [w/v] BSA, and 0.002% [v/v] Tween 20) until a response of 1 nanometer shift was reached. Loaded biosensors were dipped into kinetics buffer for 1 min to acquire a baseline and then moved to wells containing the immunogens in kinetics buffer, at 1000 nM. The trimers were allowed to associate for 180 secs before the biosensor were move back to the wells containing kinetics buffer where the baseline was acquired. Disassociation of the trimers from the IgG-loaded biosensors was recorded for 300 secs. All BLI experiments were conducted at 25°C.

### Electron Microscopy Polyclonal Epitope Mapping (EMPEM)

Plasma samples were heat-inactivated at 56°C for 1hr. Polyclonal IgG was purified from plasma using Protein A resin (GE Healthcare) and digested into Fabs using immobilized papain resin as described previously^46^. Digested Fabs were passed over Protein A resin to remove Fc and undigested IgG. BG505 SOSIP/Fab complexes were made by mixing 10-15 μg SOSIP with 500 to 1000 μg of polyclonal Fabs. The mixture was and allowed to incubate for 18 to 24 hrs at room temperature (RT). The complexes were SEC purified using a Superose^TM^ 6 Increase 10/300 GL (GE Healthcare) column to remove excess Fab prior to EM grid preparation. Fractions containing the SOSIP/Fab complexes were pooled and concentrated using 10 kDa Amicon® spin concentrators (Millipore). Samples were diluted to ∼0.03 mg/mL in TBS (0.05 M Tris pH 7.4, 0.15 M NaCl) and adsorbed onto glow discharged carbon-coated Cu400 EM grids (Electron Microscopy Sciences) and blotted after 10 seconds. The grids were then stained with 3 μL of 2% (w/v) uranyl formate, immediately blotted, and stained again for 45 secs followed by a final blot. Image collection and data processing was performed as described previously on a FEI Talos microscope (1.98 Å/pixel; 72,000× magnification) with an electron dose of ∼25 electrons/Å^2^ using Leginon^71,72^. 2D classification, 3D sorting and 3D refinement conducted using Relion v3.0^73^. EM density maps were visualized using UCSF Chimera^74^ and UCSF ChimeraX^75^ and segmented using Segger^76^.

### CryoEMPEM sample preparation

∼7 mg of clean polyclonal fab from animal Rh.32613 at week 42 time point was complexed with 200 µg of BG505 SOSIP Boost 2 antigen. Sample was allowed to complex overnight (∼18 hours) at room temperature and were SEC purified on a HiLoad 16/600 Superdex 200 pg column (GE Healthcare) using TBS as a running buffer. Sample was then concentrated to 6.1 mg/ml for application onto cryoEM grids.

### CryoEMPEM-grid preparation and CryoEMPEM imaging

Experiments were carried out as described previously.^47^ We used a Vitrobot mark IV (Thermo Fisher Scientific) for cryo-grid preparation. Chamber temperature was set to 10°C, maintained humidity at 100% with a varied blotting time within a range of 4.5-7.5 s, and blotting force was set to 0 with a wait time of 10 s. Prior to sample application, we combined the sample with lauryl maltose neopentyl glycol (LMNG) at a final concentration of 0.005 mM. We used UltrAuFoil R 1.2/1.3 (Au, 300-mesh; Quantifoil Micro Tools GmbH) grids for freezing. Grids were also treated with Ar/O_2_ (Solaris plasma cleaner, Gatan) for 10 s. 3 µl of sample with LMNG were applied to the grid before being blot and then plunge-frozen into liquid-nitrogen-cooled liquid ethane. Grids were imaged using a K3 Summit direct electron detector camera (Gatan) that was mounted to an FEI Titan Krios electron microscope (Thermo Fisher Scientific) operating at 300 kV, which also had an autoloader. Images were collected at an exposure magnification of 97,000 resulting in pixel size at the specimen plane of 0.5155 Å.

### CryoEMPEM – data processing

CryoEMPEM processing was performed similarly as described previously^47^. MotionCor2^77^ was used for micrograph movie frame alignment and dose-weighting. Data processing was initially performed using cryoSPARCv2.15^78^. We used GCTF for CTF parameter estimation^79^. From cryoSPARC, trimer-fab complex particles were picked using template picker and run through two rounds of 2D classification to remove bad particle picks. From cryoSPARC, particles were transferred to Relion v3.0 to continue processing^73^. In Relion, particles then underwent a third round of 2D classification followed by 3D refinement on selected particles. Next, we followed with a continue-refine job with a mask around the trimer only and then the map was postprocessed with the trimer-only mask. We performed a round of CTF refinement on these particles, which were then used as inputs for another round of 3D refinement. The resulting particle stack was then symmetry expanded for the next step, focused classification. A 40 Å sphere mask was appropriately placed in chimera around 6 epitopes: FP/N611, IF, V1V2V3, N355/N289, N618/N625 and the Base. We next generated trimer-Fab masks for each trimer-Fab maps resolved in the focused classification. Maps were postprocessed using the corresponding trimer-Fab mask. Two additional rounds of 3D classification were performed with tighter masks around the trimer-Fv (variable domain of the Fab). Maps were again postprocessed and then subjected to a CTF-refinement for each sorted class of particles. 3D refinement was repeated on particles after the CTF refinement step. Maps were postprocessed using a tight Trimer-Fv mask one final time to generate the high-resolution maps interpreted above (Fig S4).

### CryoEMPEM – model building and refinement

Relion postprocessed maps were used for model building and refinement. The BG505 SOSIP structure from PDB entry 6V0R^27^ was docked into each map using UCSF Chimera and was then mutated to match BG505 SOSIP Boost 2 used in this study. Polyalanine Fab models were also docked into each map. Heavy and light chains were assigned by comparing CDR3 lengths and the conformations of FR2 and FR3 between the H and L chains in the maps. CDR lengths were adjusted using manual model building in Coot^80,81^ and the entire complex was refined in Rosetta^82^ and Phenix^83^. For FP1 and FP3, each position in the heavy and light chain was mutated based on side chain density and assigned a confidence value as described in Antanasijevic et al. 2022^56^. Briefly, each amino acid position was assigned a hierarchical category identifier which takes into consideration the degree of certainty and represents a predefined subset of amino acid residues that best correspond to the side chain density. The conserved disulfide pair of each chain and other IMGT anchor residues (i.e. W41) served as a fiducial markers. Sequences were then searched in a Juypter Notebook (www.jupyter.com) environment against the Indian origin rhesus macaque Germ Line Database (GLD) ^26^. Individual scores were analyzed for FR1-3, and CDR1-2 to deduce the V_H_ and V_K_/V_L_ gene usage of the predicted sequences. Top scoring sequences were analyzed with respect to the cryoEMPEM map, and the process was performed iteratively until convergence. Final models were evaluated using MolProbity^84^ and EMRinger^85^ and deposited to the PDB with polyclonal Fv regions modeled as polyalanine.

## Data Availability

CryoEMPEM structures and densities with <4 Å were deposited to PDB and EMDB, respectively, under the following accession codes: **Base4:** 8SWX, EMD-40824; **FP1:** 8SW7, EMD-40810; **FP3:** 8T2E, EMD-40981; **IF1:** 8SWV, EMD-40822; **IF3:** 8SWW, EMD-40823; **N289:** 8T2F, EMD-40982. CryoEMPEM structures and densities >4 Å were deposition to EMDB with the following accession codes: **Base1:** EMD-40807; **Base2:** EMD-40808; **Base3:** EMD-40809; **FP2:** EMD-40803; **FP4:** EMD-40804; **V1V2V3:** EMD-40805; **N625:** EMD-40806. Negative stain densities were deposited to EMDB with the follow deposition numbers: D_1000275071-D_1000275080 and D_1000275082-D_1000275085.

## Acknowledgements

The authors thank Bill Anderson, Hannah L. Turner, Charles A. Bowman and Jean-Christophe Ducom (The Scripps Research Institute) for their help with electron microscopy, data acquisition and processing. The authors also acknowledge Lauren Holden for her help on the preparation of this manuscript. This research was supported in part by the National Institute of Allergy and Infectious Diseases R01 AI145629 and P01 AI048240 (to DJI), the National Institute of Allergy and Infectious Diseases, Center for HIV/AIDS Vaccine Immunology and Immunogen Discovery UM1AI10066, Consortium for HIV/AIDS Vaccine Development UM1 AI144462. C.A.C. was supported by a NIH F31 Ruth L. Kirschstein Predoctoral Award Al131873 and by the Achievement Rewards for College Scientists Foundation. A.A. was supported by amfAR Mathilde Krim Fellowship in Biomedical Research #110182-69-RKVA. DJI is an investigator of the Howard Hughes Medical Institute. The NHP studies were supported by the Comprehensive Antibody Vaccine Immune Consortium (CAVIMC) for National Institute of Health P51 OD011132. A portion of this research was supported by NIH grant U24GM129547 and performed at the Collaboration of AID Vaccine Discovery (CAVD) (INV-007368) as well as PNCC at OHSU and accessed through EMSL (grid.436923.9), a DOE Office of Science User Facility sponsored by the Office of Biological and Environmental Research. The contents of this manuscript are the responsibility of the authors and do not necessarily reflect the views of NIAID or the US Government. The funders had no role in study design, data collection and analysis, decision to publish or preparation of the manuscript.

## Author Contributions

C.A.C. – Conceptualization, Data curation, Formal Analysis, Investigation, Methodology, Validation, Visualization, Writing - original draft, Writing – review & editing. P.P.P – Data curation, Formal Analysis, Investigation, Visualization, Writing – original draft, Writing – review & editing. K.M.C. – Formal Analysis, Investigation, Visualization, Writing – review and editing. D.G.C. – Resources. C.A.E. – Resources. A.A. – Formal Analysis, Investigation, Methodology, Software. G.O. – Formal Analysis, Investigation, Methodology, Software, Visualization, Writing – review & editing. L.M.S. – Investigation. H.G. – Investigation, Resources. J.D.A. – Formal Analysis, Investigation, Visualization. B.N. – Investigation. M.S. – Investigation. J.B. – Investigation, Writing – review and editing. M.P. – Investigation. D.J.I – Resources. D.M. – Resources. M.C. – Resources. D.R.B – Funding acquisition, Supervision. G.S. – Resources, S.C. – Conceptualization, Funding Acquisition, Project Administration, Supervision, Writing – review and editing. A.B.W. – Conceptualization, Funding acquisition, Project Administration, Supervision, Writing – review and editing.

