## Supplemental Information for "Focusing antibody responses to the fusion peptide in rhesus macaques"

**Short Title/ Running Title: Fusion Peptide Targeting in NHPs**

Christopher A. Cottrell[1,2,#], Payal P. Pratap[1,2,#], Kimberly M. Cirelli[3], Diane G. Carnathan[2,4], Chiamaka A Enemuo[4], Aleksandar Antanasijevic[1,2], Gabriel Ozorowski[1,2], Leigh M. Sewall[1,2], Hongmei Gao[5], Joel D. Allen[6], Bartek Nogal[1,2], Murillo Silva[8], Jinal Bhiman[9], Matthias Pauthner[10], Darrell J. Irvine[2,8], David Montefiori[5], Max Crispin[6], Dennis R. Burton[2,7,10], Guido Silvestri[2,4], Shane Crotty[2,3,11], and Andrew B. Ward[1,2]*

**Author Affiliations**

[1] Department of Integrative Structural and Computational Biology, The Scripps Research Institute, La Jolla, CA 92037, USA

[2]International AIDS Vaccine Initiative Neutralizing Antibody Center, the Collaboration for AIDS Vaccine Discovery (CAVD) and Scripps Consortium for HIV/AIDS Vaccine Development (CHAVD), The Scripps Research Institute, La Jolla, CA 92037, USA.

[3]La Jolla Institute for Immunology, La Jolla, CA 92037, USA.

[4]Division of Microbiology and Immunology, Yerkes National Primate Research Center, Emory University, Atlanta, GA 30329, USA.

[5]Duke Human Vaccine Institute and Department of Surgery, Duke University Medical Center Durham, NC, USA.

[6]School of Biological Sciences, University of Southampton, Southampton, SO17 1BJ, UK

[7]Department of Immunology and Microbiology, The Scripps Research Institute, La Jolla, California, USA.

[8]Koch Institute for Integrative Cancer Research, Massachusetts Institute of Technology, Cambridge, MA 02139, USA.

[9]Centre for HIV and STI, National Institute for Communicable Diseases of the National Health Laboratory Service, Johannesburg, South Africa.

[10]Center for HIV/AIDS Vaccine Immunology and Immunogen Discovery and IAVI Neutralizing Antibody Center, Department of Immunology and Microbiology, The Scripps Research Institute, La Jolla, California, USA.

[11]Division of Infectious Disease and Global Public Health, Department of Medicine, University of California, San Diego, La Jolla, California, USA

[#]Contributed equally

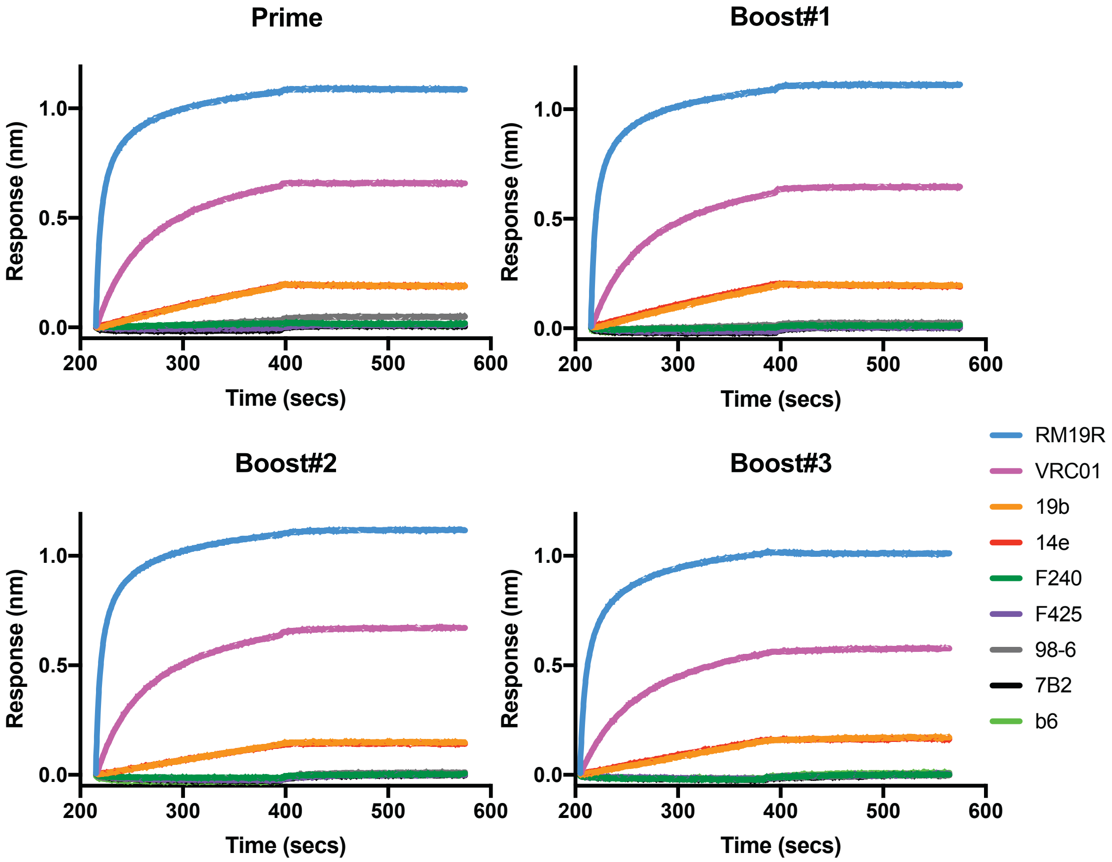

**Figure S1. Antigenicity of FP targeting immunogen series.** RM19R is a base-binding mAb elicited in a BG505 SOSIP.664 immunized RM (*9*). VRC01 is a CD4 binding site targeting bnAb (*34*) and b6 is a CD4 binding site targeting non-neutralizing mAb. 19b, 14e, and F425 are V3 targeting non-neutralizing mAbs. F240, 7B2, and 98-6 post-fusion gp41 targeting mAbs.

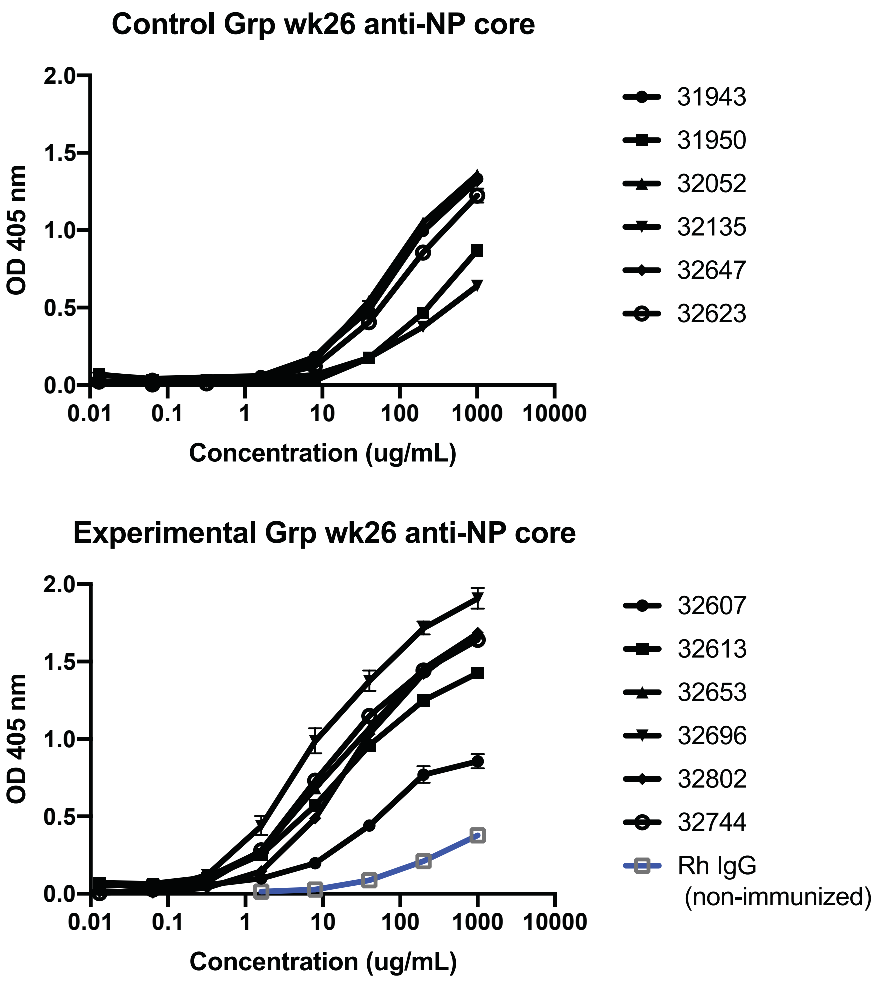

**Fig. S2. IgG binding ELISAs for T33-31 nanoparticle core.**

IgG purified from week 26 was analyzed at 5-fold dilutions to determine IgG recognition of T33-31 nanoparticle core. Upper panel shows response for the control group monkeys while lower panel shows response for the experimental group monkeys.

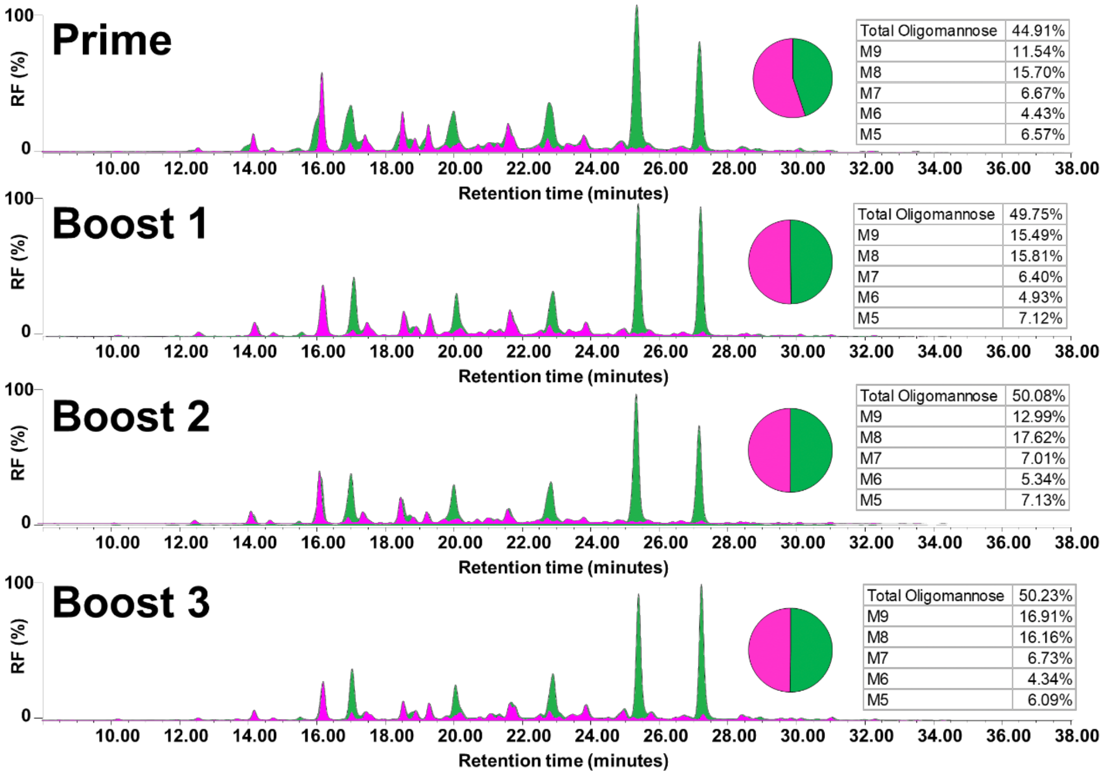

**Fig. S3. UPLC analysis of released N-glycans from the sequential immunogens.** Procainamide labelled glycans were subjected to endoH digestion that enables the determination and quantification of oligomannose and hybrid-type glycans (green) and complex-type glycans (magenta). Oligomannose-type glycans were quantified using the Waters Empower software.

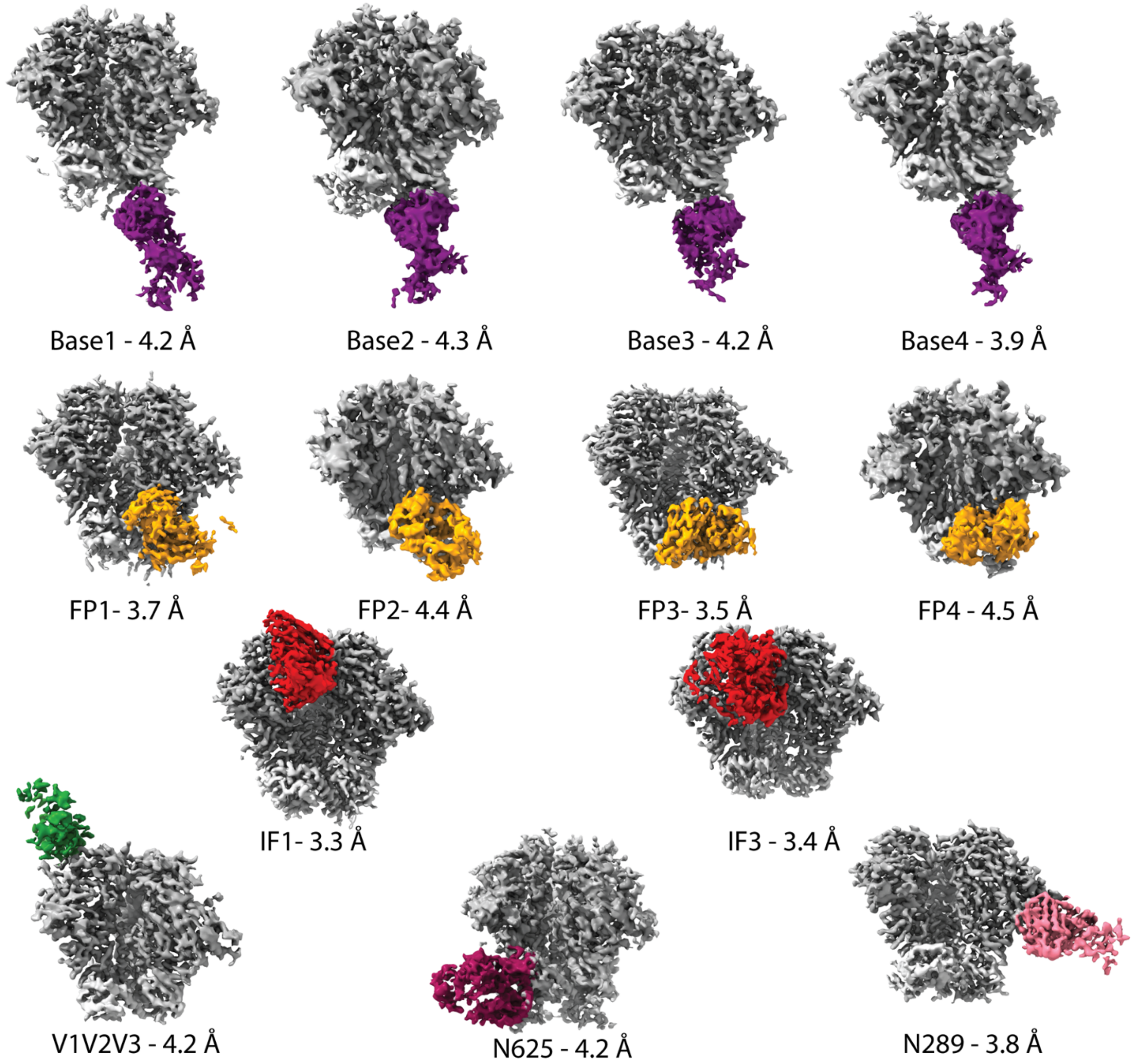

**Fig S4. CryoEMPEM High Resolution Maps**

Thirteen high resolution (≤4.5 Å) maps were resolved from one polyclonal sample (animal 32613 at week 42 timepoint). Resolved maps include four against the base of the trimer, four maps against the fusion peptide (FP) region, two against the interface (IF) region, one against the V1V2V3 variable region, one against the N625 gp41 glycan and one against N289 glycan of gp120.

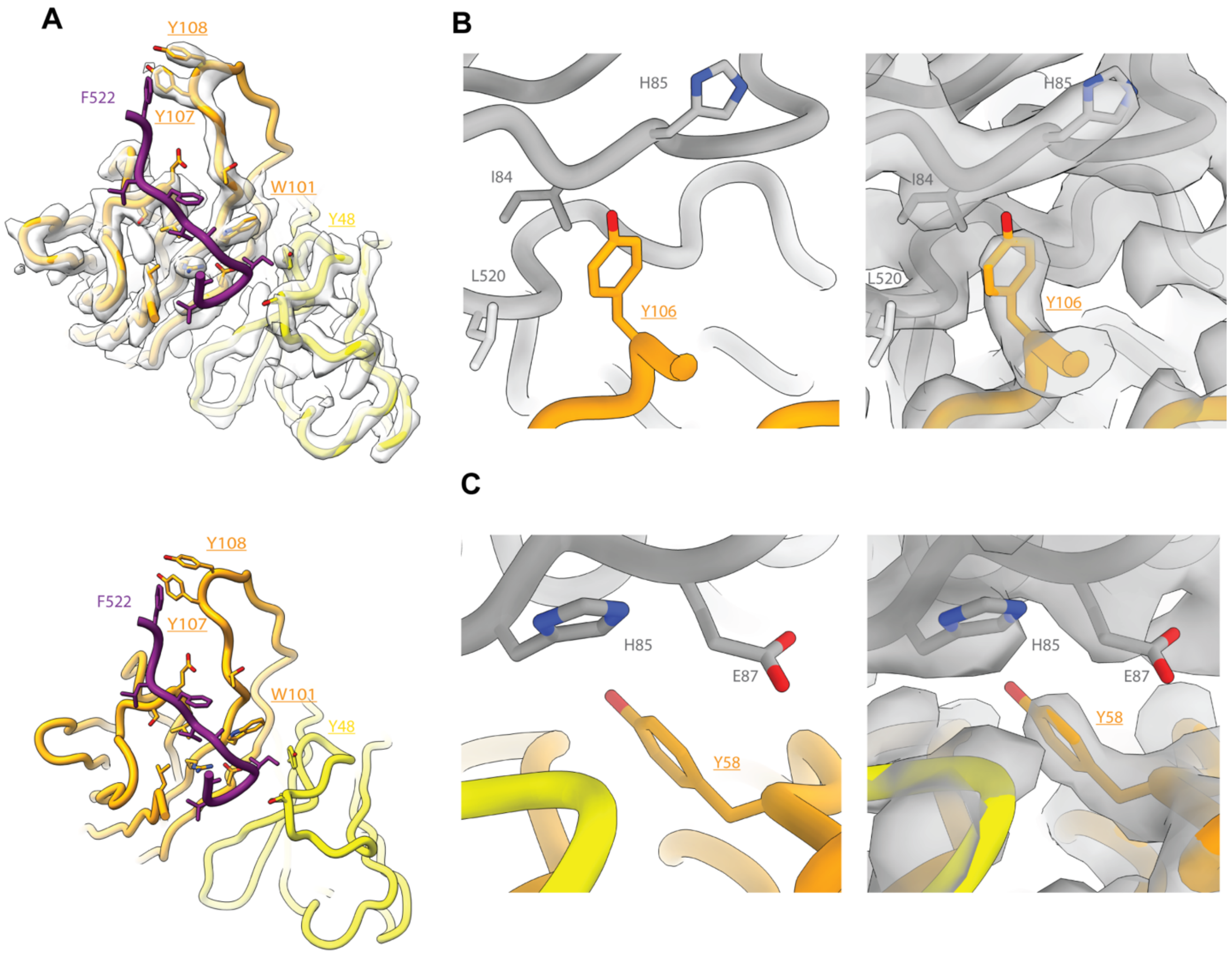

Figure S5. CryoEMPEM Polyclonal Residue Assignments

A) FP3 map and model showing residue predictions (underlined residues) based on density information for FP interaction with HCDR3 and LFW2 aromatic residues. B) FP1 map and model showing residue predictions (underlined residues) based on density information for HCDR3 interaction with C-terminal FP residues and C1 region of gp120. C) FP1 map and model showing residue prediction (underlined residue) based on density information for HCDR2 interaction with C1 region of gp120.

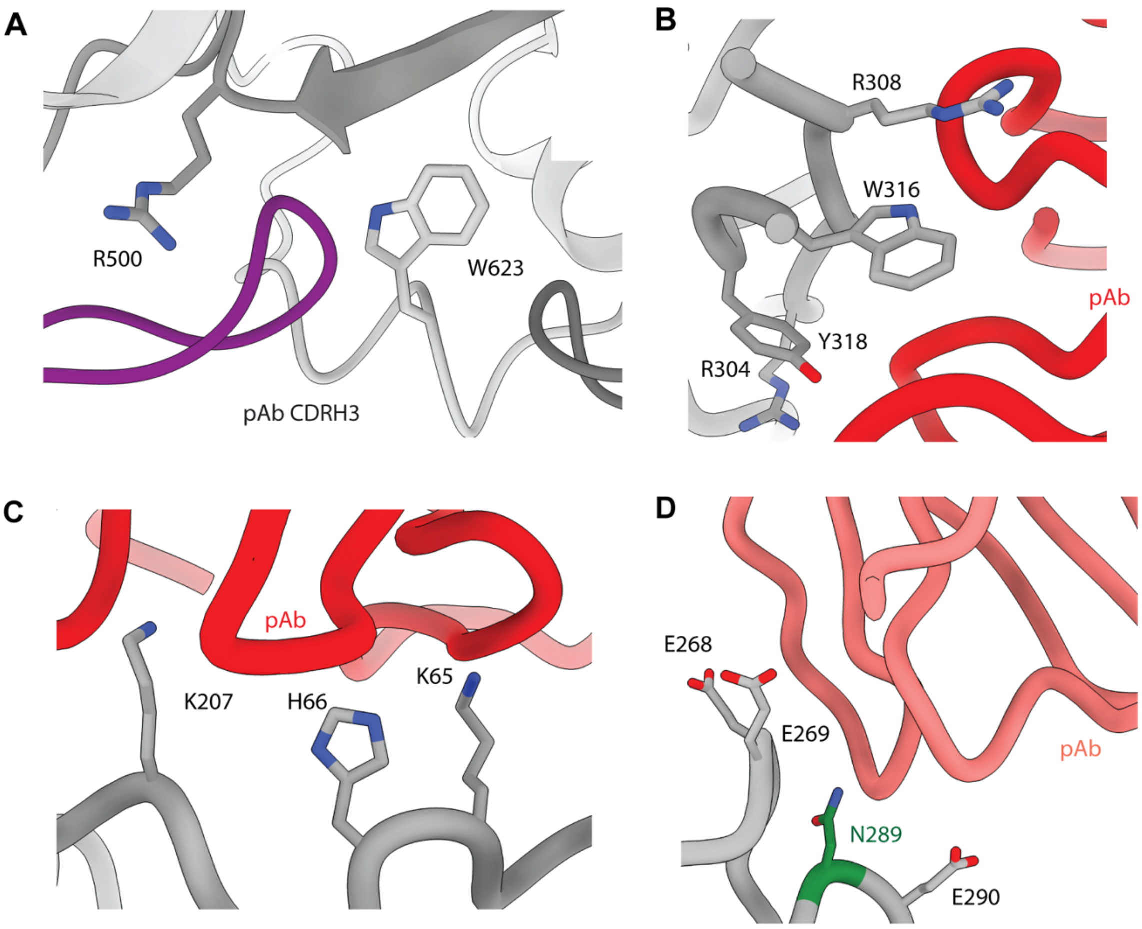

Figure S6. CryoEM Analysis of Off-target Responses

A) Base response elucidated from Base-4 map shows predicted CDRH3 of pAb interaction with base of the trimer. B) IF response elucidated from IF-1 map shows antibody interactions with V3 region of trimer. C) IF response elucidated from IF-3 map shows antibody interactions with C1 region of trimer. D) N289 response shows antibody interaction with C2 region of trimer due to absence of N289 glycan.

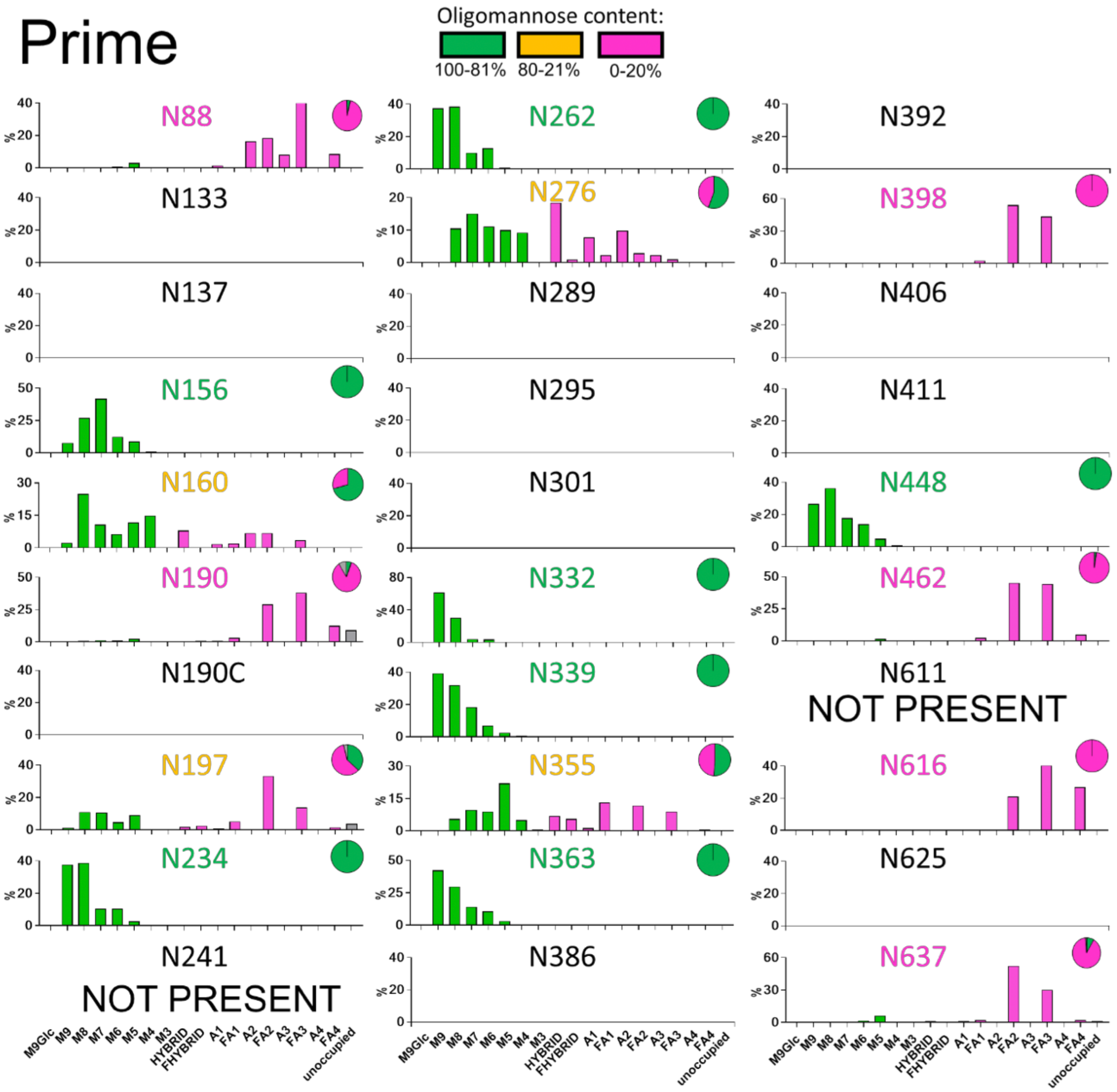

**Fig. S7. Site-specific analysis of the Prime immunogen.**

The intensity of different glycoforms at a single PNGS were compared and glycopeptides were categorized according to the number of mannose residues (green), hybrids that possess one unprocessed arm and one processed and also the number of processed branches and the presence/absence of fucose for complex-type glycans (magenta). The proportion of unoccupied asparagines at each site is colored grey. Pie charts summing the total oligomannose (M9-M4), complex and unoccupied are shown to the right of each site and the color of the lettering refers to the % oligomannose at each site, with green letters containing over 80% oligomannose, between 80% and 20% colored orange and less than 20% magenta.

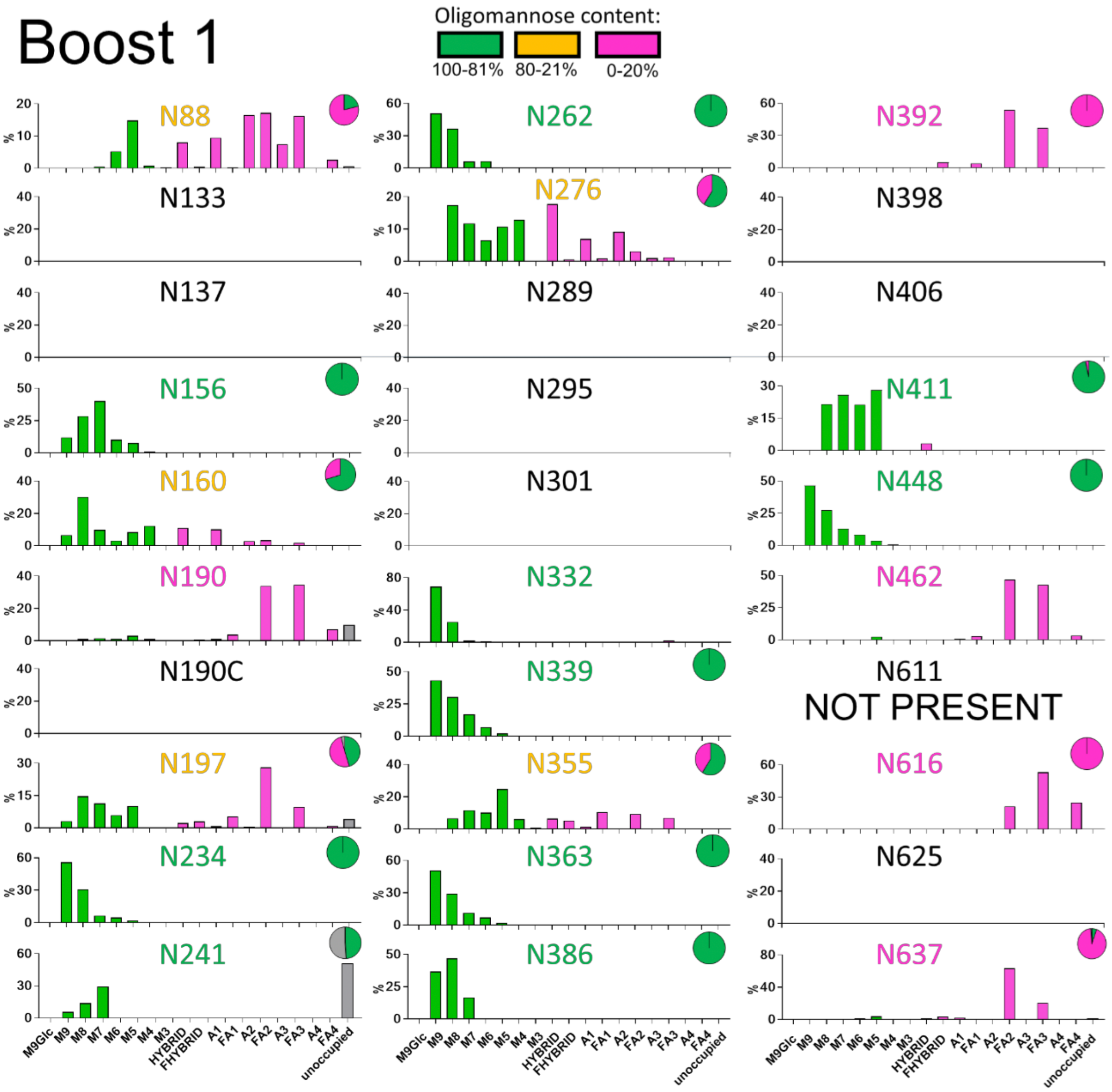

**Fig. S8. Site-specific analysis of the Boost#1 immunogen.**

Same as in Fig. S7

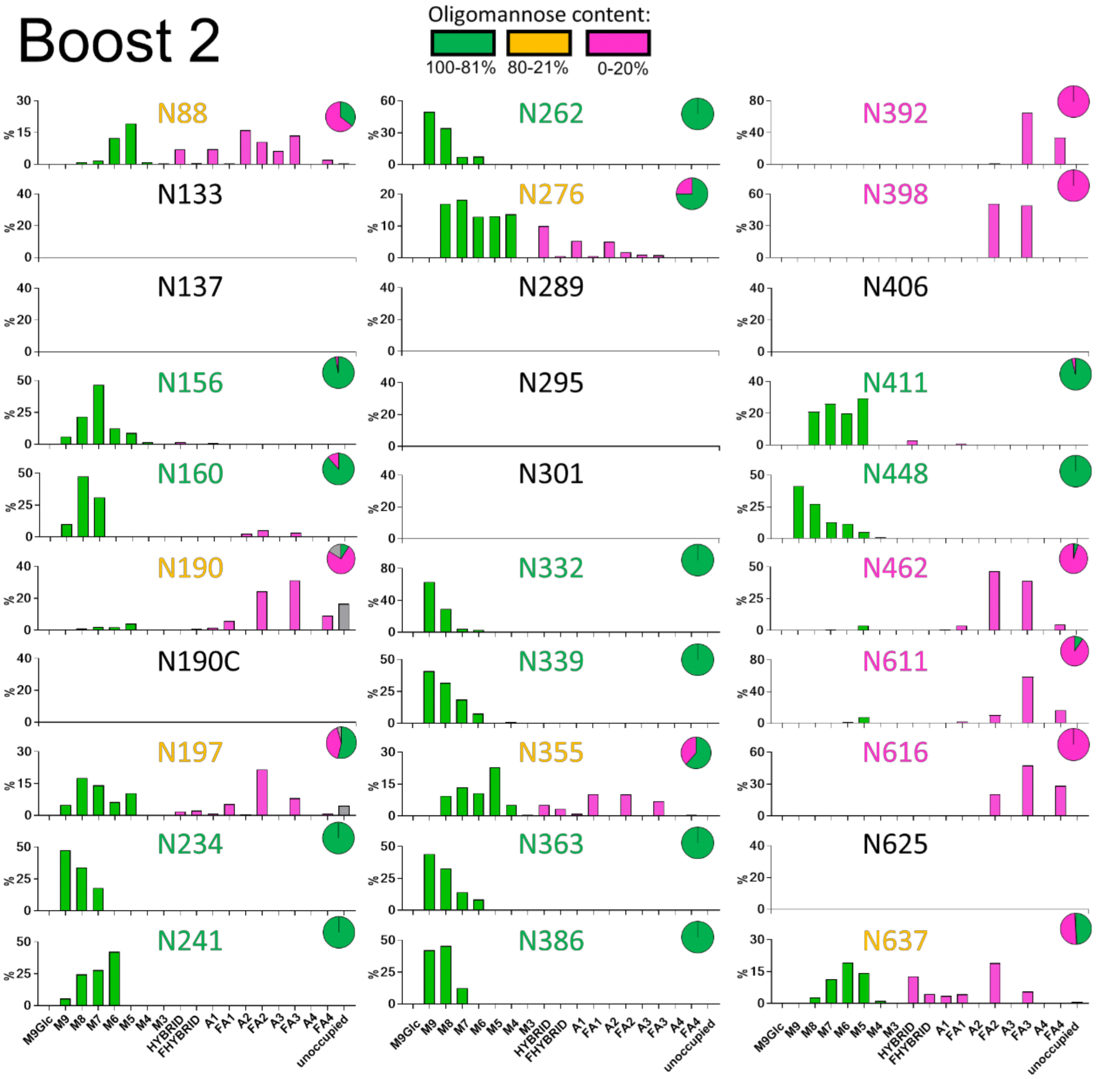

**Fig. S9. Site-specific analysis of the Boost#2 immunogen.**

Same as in Fig. S7

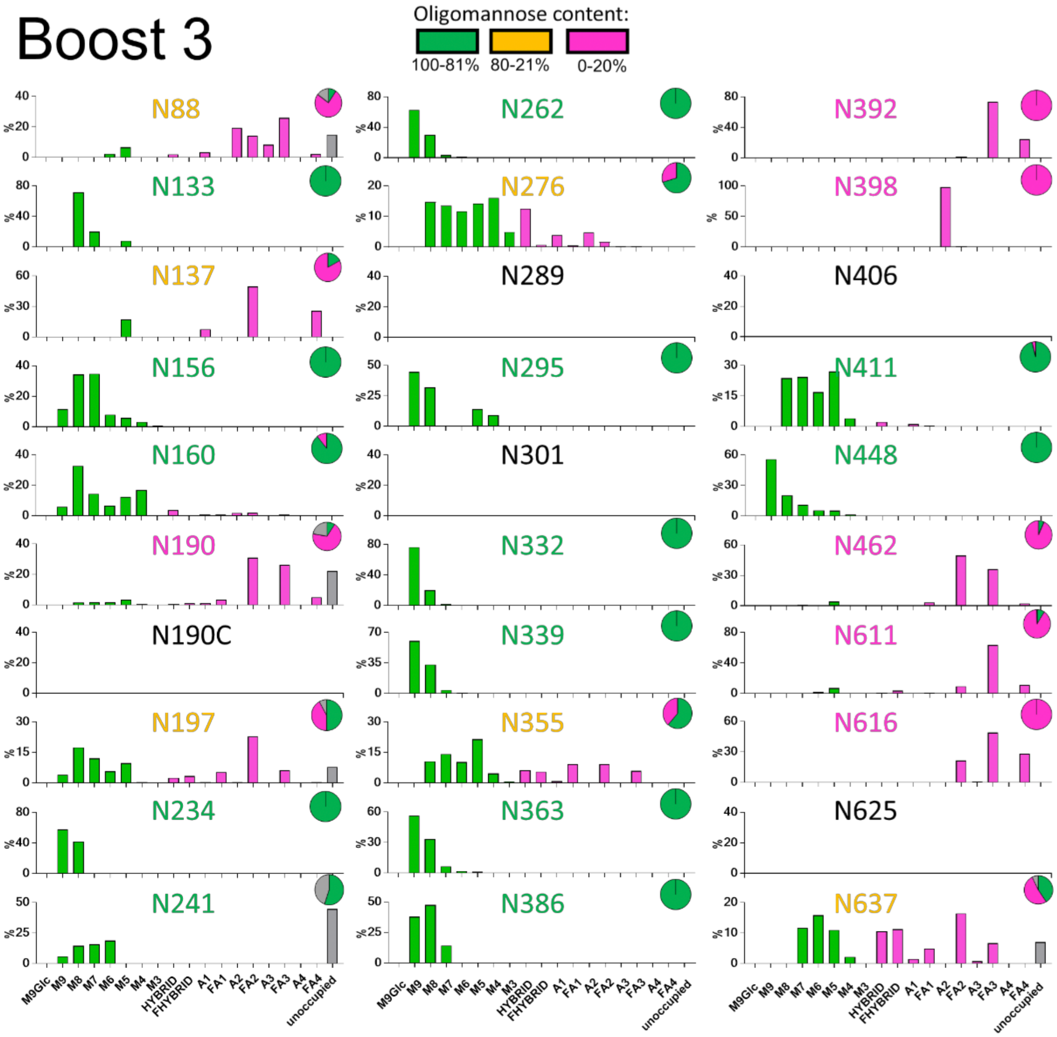

**Fig. S10. Site-specific analysis of the Boost#3 immunogen.**

Same as in Fig. S7

**Table S1. CryoEM data collection, processing and model building statistics.**

| Map | BG505_BOOST_2 32613 WK 42 Polyclonal Fab Base1 | BG505_BOOST_2 32613 WK 42 Polyclonal Fab Base2 | BG505_BOOST_2 32613 WK 42 Polyclonal Fab Base3 | BG505_BOOST_2 32613 WK 42 Polyclonal Fab Base4 |
|---|---|---|---|---|
| EMDB | EMD-40807 | EMD-40808 | EMD-40809 | EMD-40824 |
| **Data collection** | | | | |
| Microscope | TFS Titan Krios | TFS Titan Krios | TFS Titan Krios | TFS Titan Krios |
| Voltage (kV) | 300 | 300 | 300 | 300 |
| Detector | Gatan K3 | Gatan K3 | Gatan K3 | Gatan K3 |
| Recording mode | Super-resolution | Super-resolution | Super-resolution | Super-resolution |
| Nominal magnification (without super-res) | 22,500x | 22,500x | 22,500x | 22,500x |
| Movie micrograph pixelsize (Å; super-res) | 0.5155 | 0.5155 | 0.5155 | 0.5155 |
| Number of frames | 71 | 71 | 71 | 71 |
| Total dose ($e^-/Å^2$) | 40.9 | 40.9 | 40.9 | 40.9 |
| Defocus range (µm) | -2.0 to -0.5 | -2.0 to -0.5 | -2.0 to -0.5 | -2.0 to -0.5 |
| **EM data processing** | | | | |
| Number of movie micrographs | 6015 | 6015 | 6015 | 6015 |
| Number of molecular projection images in map | 56597 | 44865 | 47856 | 53702 |
| Symmetry | C1 | C1 | C1 | C1 |
| Map pixel size | 1.031 | 1.031 | 1.031 | 1.031 |
| Map resolution (FSC 0.143; Å) | 4.2 | 4.3 | 4.2 | 3.9 |
| Map sharpening B-factor ($Å^2$) | -111.11 | -130.81 | -103.58 | -98.19 |
| **Structure building and validation** | | | | |
| *Number of atoms in deposited model* | | | | |
| Non-hydrogen atoms | *NA* | *NA* | *NA* | 15170 |
| Protein residues | | | | 1866 |
| ligands | | | | 67 |
| MolProbity score | | | | 1.10 |
| Clashscore | | | | 1.27 |
| EMRinger score | | | | 2.83 |
| d FSC model (0.5; Å) | | | | 4.0 |
| *RMSD from ideal* | | | | |
| Bond length (Å) | | | | 0.021 |
| Bond angles (˚) | | | | 1.678 |
| *Ramachandran plot* | | | | |
| Favored (%) | | | | 96.26 |
| Allowed (%) | | | | 3.41 |
| Outliers (%) | | | | 0.33 |
| Side chain rotamer outliers (%) | | | | 0.41 |
| Cβ outliers (%) | | | | 0.00 |
| PDB | | | | 8SWX |

| Map | BG505_BOOST_2 32613 WK 42 Polyclonal Fab FP1 | BG505_BOOST_2 32613 WK 42 Polyclonal Fab FP2 | BG505_BOOST_2 32613 WK 42 Polyclonal Fab FP3 | BG505_BOOST_2 32613 WK 42 Polyclonal Fab FP4 |
|---|---|---|---|---|
| EMDB | EMD-40810 | EMD-40803 | EMD-40981 | EMD-40804 |
| **Data collection** | | | | |
| Microscope | TFS Titan Krios | TFS Titan Krios | TFS Titan Krios | TFS Titan Krios |
| Voltage (kV) | 300 | 300 | 300 | 300 |
| Detector | Gatan K3 | Gatan K3 | Gatan K3 | Gatan K3 |
| Recording mode | Super-resolution | Super-resolution | Super-resolution | Super-resolution |
| Nominal magnification | 22,500x | 22,500x | 22,500x | 22,500x |
| Movie micrograph pixelsize (Å) | 0.5155 | 0.5155 | 0.5155 | 0.5155 |
| Number of frames (Falcon 4 EER fractions) | 71 | 71 | 71 | 71 |
| Total dose ($e^-/Å^2$) | 40.9 | 40.9 | 40.9 | 40.9 |
| Defocus range (µm) | -2.0 to -0.5 | -2.0 to -0.5 | -2.0 to -0.5 | -2.0 to -0.5 |
| **EM data processing** | | | | |
| Number of movie micrographs | 6015 | 6015 | 6015 | 6015 |
| Number of molecular projection images in map | 63946 | 40085 | 88235 | 31418 |
| Symmetry | C1 | C1 | C1 | C1 |
| Map pixel size | 1.031 | 1.031 | 1.031 | 1.031 |
| Map resolution (FSC 0.143; Å) | 3.7 | 4.3 | 3.5 | 4.4 |
| Map sharpening B-factor ($Å^2$) | -102.76 | -134.84 | -96.84 | -103.84 |
| **Structure building and validation** | | | | |
| *Model Composition* | | | | |
| Non-hydrogen atoms | 15147 | *NA* | 14900 | *NA* |
| Protein residues | 1895 | | 1852 | |
| ligands | 59 | | 64 | |
| MolProbity score | 0.96 | | 0.71 | |
| Clashscore | 0.97 | | 0.64 | |
| EMRinger score | 2.95 | | 3.20 | |
| d FSC model (0.5; Å) | 3.9 | | 3.6 | |
| *RMSD from ideal* | | | | |
| Bond length (Å) | 0.022 | | 0.021 | |
| Bond angles (˚) | 1.706 | | 1.625 | |
| *Ramachandran plot* | | | | |
| Favored (%) | 97.03 | | 97.99 | |
| Allowed (%) | 2.76 | | 1.96 | |
| Outliers (%) | 0.22 | | 0.06 | |
| Side chain rotamer outliers (%) | 0.14 | | 0.28 | |
| Cβ outliers (%) | 0.00 | | 0.00 | |
| PDB | 8SW7 | | 8T2E | |

| Map | BG505_BOOST_2 32613 WK 42 Polyclonal Fab IF1 | BG505_BOOST_2 32613 WK 42 Polyclonal Fab IF3 | BG505_BOOST_2 32613 WK 42 Polyclonal Fab V1V2V3 | BG505_BOOST_2 32613 WK 42 Polyclonal Fab N625 |
|---|---|---|---|---|
| EMDB | EMD-40822 | EMD-40823 | EMD-40805 | EMD-40806 |
| **Data collection** | | | | |
| Microscope | TFS Titan Krios | TFS Titan Krios | TFS Titan Krios | TFS Titan Krios |
| Voltage (kV) | 300 | 300 | 300 | 300 |
| Detector | Gatan K3 | Gatan K3 | Gatan K3 | Gatan K3 |
| Recording mode | Super-resolution | Super-resolution | Super-resolution | Super-resolution |
| Nominal magnification (without super-res) | 22,500x | 22,500x | 22,500x | 22,500x |
| Movie micrograph pixelsize (Å; super-res) | 0.5155 | 0.5155 | 0.5155 | 0.5155 |
| Number of frames | 71 | 71 | 71 | 71 |
| Total dose ($e^-/Å^2$) | 40.9 | 40.9 | 40.9 | 40.9 |
| Defocus range (μm) | -2.0 to -0.5 | -2.0 to -0.5 | -2.0 to -0.5 | -2.0 to -0.5 |
| **EM data processing** | | | | |
| Number of movie micrographs | 6015 | 6015 | 6015 | 6015 |
| Number of molecular projection images in map | 231106 | 99215 | 54747 | 38218 |
| Symmetry | C1 | C1 | C1 | C1 |
| Map pixel size | 1.031 | 1.031 | 1.031 | 1.031 |
| Map resolution (FSC 0.143; Å) | 3.3 | 3.4 | 4.2 | 4.2 |
| Map sharpening B-factor ($Å^2$) | -103.59 | -95.39 | -134.42 | -113.30 |
| **Structure building and validation** | | | *NA* | |
| *Number of atoms in deposited model* | | | | |
| Non-hydrogen atoms | 14906 | 14840 | | |
| Protein residues | 1853 | 1881 | | |
| ligands | 60 | 43 | | |
| MolProbity score | 0.93 | 0.95 | | |
| Clashscore | 0.81 | 0.85 | | |
| EMRinger score | 3.83 | 3.85 | | |
| d FSC model (0.5; Å) | 3.4 | 3.6 | | |
| *RMSD from ideal* | | | | |
| Bond length (Å) | 0.021 | 0.021 | | |
| Bond angles (°) | 1.635 | 1.679 | | |
| *Ramachandran plot* | | | | |
| Favored (%) | 96.99 | 96.85 | | |
| Allowed (%) | 2.89 | 3.15 | | |
| Outliers (%) | 0.11 | 0.00 | | |
| Side chain rotamer outliers (%) | 0.21 | 0.00 | | |
| Cβ outliers (%) | 0.00 | 0.00 | | |
| PDB | 8SWV | 8SWW | | |

| **Map** | **BG505_BOOST_2 32613 WK 42 Polyclonal Fab N289** |
|---|---|
| EMDB | EMD-40982 |
| **Data collection** | |
| Microscope | TFS Titan Krios |
| Voltage (kV) | 300 |
| Detector | Gatan K3 |
| Recording mode | Super-resolution |
| Nominal magnification (without super-res) | 22,500x |
| Movie micrograph pixelsize (Å; super-res) | 0.5155 |
| Number of frames | 71 |
| Total dose ($e^-$/Å$^2$) | 40.9 |
| Defocus range (µm) | -2.0 to -0.5 |
| **EM data processing** | |
| Number of movie micrographs | 6015 |
| Number of molecular projection images in map | 75994 |
| Symmetry | C1 |
| Map pixel size | 1.031 |
| Map resolution (FSC 0.143; Å) | 3.8 |
| Map sharpening B-factor (Å$^2$) | -115.80 |
| **Structure building and validation** | |
| *Number of atoms in deposited model* | |
| Non-hydrogen atoms | 14097 |
| Protein residues | 1762 |
| ligands | 56 |
| MolProbity score | 0.96 |
| Clashscore | 0.72 |
| EMRinger score | 2.58 |
| d FSC model (0.5; Å) | 4.0 |
| *RMSD from ideal* | |
| Bond length (Å) | 0.021 |
| Bond angles (˚) | 1.678 |
| *Ramachandran plot* | |
| Favored (%) | 96.52 |
| Allowed (%) | 3.42 |
| Outliers (%) | 0.06 |
| Side chain rotamer outliers (%) | 0.22 |
| Cβ outliers (%) | 0.00 |
| PDB | 8T2F |

**Table S2. BG505 pseudovirus neutralization data weeks 24, 26 and 42.**

| Study Group | Animal ID | Week | SVA-MLV | BG505 | BG505/ T332N | BG505/ T332N. N611A | BG505/T3 32N. T465N | BG505/ T332N. 133aN+ 136aA |
|---|---|---|---|---|---|---|---|---|
| Control | 31943 | 24 | <20 | **2453** | **1409** | **1383** | **621** | **409** |
| Control | 31950 | 24 | <20 | **202** | **217** | **323** | **60** | **240** |
| Control | 32052 | 24 | <20 | **613** | **866** | **836** | **68** | **714** |
| Control | 32135 | 24 | <20 | **318** | **290** | **333** | <20 | **347** |
| Control | 32647 | 24 | <20 | <20 | <20 | <20 | <20 | <20 |
| Control | 32623 | 24 | <20 | **419** | **584** | **804** | **75** | **580** |
| Experimental | 32607 | 24 | <20 | **356** | **200** | **398** | **50** | **57** |
| Experimental | 32613 | 24 | <20 | <20 | <20 | **282** | <20 | <20 |
| Experimental | 32653 | 24 | <20 | 22 | 22 | **227** | <20 | 28 |
| Experimental | 32696 | 24 | <20 | 24 | **74** | **558** | **46** | 24 |
| Experimental | 32802 | 24 | <20 | <20 | 22 | **676** | <20 | <20 |
| Experimental | 32744 | 24 | <20 | 23 | <20 | **174** | <20 | <20 |
| Control | 31943 | 26 | <20 | **2069** | **1098** | **1000** | **643** | **199** |
| Control | 31950 | 26 | <20 | **104** | **134** | **159** | **50** | **137** |
| Control | 32052 | 26 | <20 | **305** | **314** | **412** | 28 | **365** |
| Control | 32135 | 26 | <20 | **197** | **174** | **210** | <20 | **231** |
| Control | 32647 | 26 | <20 | <20 | 28 | <20 | <20 | <20 |
| Control | 32623 | 26 | <20 | **273** | **288** | **527** | **57** | **325** |
| Experimental | 32607 | 26 | <20 | **486** | **312** | **483** | **125** | **77** |
| Experimental | 32613 | 26 | <20 | <20 | 21 | **256** | <20 | <20 |
| Experimental | 32653 | 26 | <20 | 39 | 33 | **183** | 21 | **50** |
| Experimental | 32696 | 26 | <20 | **40** | **91** | **468** | **47** | **50** |
| Experimental | 32802 | 26 | <20 | <20 | 22 | **465** | <20 | <20 |
| Experimental | 32744 | 26 | <20 | <20 | <20 | **94** | <20 | 22 |
| Control | 31943 | 42 | <20 | **1,663** | **1,134** | **1,209** | **394** | **393** |
| Control | 31950 | 42 | <20 | **1,179** | **1,199** | **1,269** | **397** | **1,190** |
| Control | 32052 | 42 | <20 | **315** | **302** | **281** | <20 | **322** |
| Control | 32135 | 42 | <20 | **4,286** | **4,366** | **5,063** | <20 | **5,070** |
| Control | 32647 | 42 | <20 | **49** | **88** | **73** | <20 | **74** |
| Control | 32623 | 42 | <20 | **1,743** | **1,737** | **1,970** | **521** | **1,777** |
| Experimental | 32607 | 42 | <20 | **1,823** | **1,361** | **2,288** | **314** | **638** |
| Experimental | 32613 | 42 | <20 | **81** | **68** | **3,589** | **47** | **50** |
| Experimental | 32653 | 42 | <20 | **125** | **80** | **534** | <20 | <20 |
| Experimental | 32696 | 42 | <20 | **224** | **234** | **643** | **42** | **37** |
| Experimental | 32802 | 42 | <20 | **101** | **98** | **4,356** | **48** | **82** |
| Experimental | 32744 | 42 | <20 | **78** | **81** | **475** | **44** | **70** |

Values are the serum dilution at which relative luminescence units (RLUs) were reduced 50% compared to virus control wells (no test sample)

**Table S3. FP-sensitive pseudovirus panel neutralization data weeks 24, 26, and 42.**

| Study Group# | Animal ID | Week | 25710-2.43 | 3988.25 | 0077.V1.C16 | CNE19 | CNE56 | KER2008 .vrc12 | Q23.17 | 286_36 | BL01.DG |
|---|---|---|---|---|---|---|---|---|---|---|---|
| Control | 31943 | 24 | <20 | <20 | <20 | **214** | <20 | <20 | <20 | <20 | <20 |
| Control | 31950 | 24 | <20 | 26 | <20 | 24 | <20 | <20 | <20 | <20 | <20 |
| Control | 32052 | 24 | <20 | 26 | <20 | <20 | <20 | <20 | <20 | <20 | <20 |
| Control | 32135 | 24 | <20 | 32 | <20 | 34 | <20 | <20 | <20 | 24 | <20 |
| Control | 32647 | 24 | <20 | 31 | <20 | <20 | <20 | <20 | <20 | <20 | <20 |
| Control | 32623 | 24 | <20 | <20 | <20 | <20 | <20 | <20 | <20 | <20 | <20 |
| Experimental | 32607 | 24 | <20 | <20 | <20 | <20 | <20 | <20 | <20 | <20 | <20 |
| Experimental | 32613 | 24 | <20 | <20 | <20 | 32 | <20 | <20 | <20 | <20 | <20 |
| Experimental | 32653 | 24 | <20 | 34 | <20 | <20 | <20 | <20 | <20 | <20 | <20 |
| Experimental | 32696 | 24 | <20 | 24 | <20 | 20 | <20 | <20 | <20 | <20 | <20 |
| Experimental | 32802 | 24 | <20 | <20 | <20 | <20 | <20 | <20 | <20 | <20 | <20 |
| Experimental | 32744 | 24 | <20 | 26 | <20 | <20 | <20 | <20 | <20 | <20 | <20 |
| Control | 31943 | 26 | <20 | <20 | <20 | **136** | <20 | <20 | <20 | <20 | <20 |
| Control | 31950 | 26 | <20 | 25 | <20 | 27 | <20 | <20 | <20 | <20 | <20 |
| Control | 32052 | 26 | <20 | <20 | <20 | <20 | <20 | <20 | <20 | <20 | <20 |
| Control | 32135 | 26 | <20 | 31 | <20 | 21 | <20 | <20 | <20 | 21 | <20 |
| Control | 32647 | 26 | <20 | **55** | <20 | 22 | <20 | <20 | <20 | <20 | 36 |
| Control | 32623 | 26 | <20 | <20 | <20 | <20 | <20 | <20 | <20 | <20 | <20 |
| Experimental | 32607 | 26 | <20 | 22 | <20 | <20 | <20 | <20 | <20 | <20 | <20 |
| Experimental | 32613 | 26 | <20 | 21 | <20 | 26 | <20 | <20 | <20 | <20 | <20 |
| Experimental | 32653 | 26 | <20 | **42** | <20 | 32 | <20 | <20 | 21 | 23 | <20 |
| Experimental | 32696 | 26 | 22 | 39 | <20 | 30 | <20 | <20 | 27 | <20 | **41** |
| Experimental | 32802 | 26 | <20 | <20 | <20 | <20 | <20 | <20 | <20 | <20 | <20 |
| Experimental | 32744 | 26 | <20 | 29 | <20 | 22 | <20 | <20 | <20 | <20 | <20 |
| Control | 31943 | 42 | <20 | <20 | <20 | <20 | <20 | <20 | <20 | <20 | <20 |
| Control | 31950 | 42 | <20 | <20 | <20 | <20 | <20 | <20 | <20 | <20 | <20 |
| Control | 32052 | 42 | <20 | <20 | <20 | <20 | <20 | <20 | <20 | <20 | <20 |
| Control | 32135 | 42 | <20 | <20 | <20 | <20 | <20 | <20 | <20 | <20 | <20 |
| Control | 32647 | 42 | <20 | <20 | <20 | <20 | <20 | <20 | <20 | <20 | <20 |
| Control | 32623 | 42 | <20 | <20 | <20 | <20 | <20 | <20 | <20 | <20 | <20 |
| Experimental | 32607 | 42 | <20 | <20 | <20 | <20 | <20 | <20 | <20 | <20 | <20 |
| Experimental | 32613 | 42 | **53** | **42** | <20 | **148** | <20 | <20 | <20 | <20 | <20 |
| Experimental | 32653 | 42 | **<20** | **27** | <20 | **<20** | <20 | <20 | <20 | <20 | <20 |
| Experimental | 32696 | 42 | <20 | <20 | nd | <20 | nd | <20 | <20 | nd | <20 |
| Experimental | 32802 | 42 | <20 | <20 | <20 | **34** | <20 | <20 | <20 | <20 | <20 |
| Experimental | 32744 | 42 | **30** | **32** | <20 | **37** | <20 | <20 | <20 | <20 | <20 |

Values are the serum dilution at which relative luminescence units (RLUs) were reduced 50% compared to virus control wells (no test sample)

**Table S4. Global panel pseudovirus neutralization data.**

| | | | ID50 (dilution) in TZM-bl cells | | | | | | | | | |
|---|---|---|---|---|---|---|---|---|---|---|---|---|
| **Study Group#** | **Animal ID** | **Week** | **CEO217** | **CE1176** | **25710** | **CH119** | **X2278** | **TRO11** | **BJOX2000** | **CNE55** | **X1632** | **246F3** |
| Crtl | 31943 | 42 | <10 | <10 | <10 | <10 | <10 | <10 | <10 | <10 | <10 | <10 |
| Crtl | 31950 | 42 | <10 | <10 | <10 | <10 | <10 | <10 | <10 | <10 | <10 | <10 |
| Crtl | 32052 | 42 | <10 | <10 | <10 | <10 | <10 | <10 | <10 | <10 | <10 | <10 |
| Crtl | 32135 | 42 | <10 | <10 | <10 | <10 | <10 | <10 | <10 | <10 | <10 | <10 |
| Crtl | 32647 | 42 | <10 | <10 | <10 | 13 | <10 | <10 | <10 | <10 | <10 | <10 |
| Crtl | 32623 | 42 | <10 | <10 | <10 | <10 | <10 | <10 | <10 | <10 | <10 | <10 |
| Expt | 32607 | 42 | <10 | <10 | 59 | 19 | <10 | <10 | <10 | <10 | <10 | <10 |
| Expt | 32613 | 42 | <10 | 13 | 13 | <10 | <10 | <10 | <10 | <10 | <10 | <10 |
| Expt | 32653 | 42 | <10 | <10 | <10 | <10 | <10 | <10 | <10 | <10 | <10 | <10 |
| Expt | 32696 | 42 | <10 | <10 | <10 | <10 | <10 | <10 | <10 | <10 | <10 | <10 |
| Expt | 32802 | 42 | <10 | <10 | 15 | <10 | <10 | <10 | <10 | <10 | 12 | <10 |
| Expt | 32744 | 42 | <10 | 13 | 75 | <10 | <10 | <10 | <10 | <10 | 15 | <10 |

Values are the serum dilution at which relative luminescence units (RLUs) were reduced 50% compared to virus control wells (no test sample)

**Table S5. Immunogen mutations relative to BG505 SOSIP.v5.2**

| Immunogen | Mutations relative to BG505 SOSIP.v5.2 |
|---|---|
| Prime | M271I-F288L-T290E-P291S-N611Q |
| Boost#1 | P240T-S241N-M271I-F288L-T290E-P291S-N611Q |
| Boost#2 | P240T-S241N-M271I-F288L-T290E-P291S-S613T |
| Boost#3 | H85V-K229N-P240T-S241N-M271I-F288L-T290E-P291S-S613T |

**Table S6. FACS antibody panel.**

| Marker | Clone | Flurophore |
|---|---|---|
| Viability | n/a | APCe780 |
| CD20 | 2H7 | Ax488 |
| CD4 | OKT-4 | APCe780 |
| CD8a | RPA-T8 | APCe780 |
| CD16 | ebioCB16 | APCe780 |
| IgG | G18-145 | AF700 |
| IgM | G20-127 | PerCP-Cy5.5 |
| CD71 | L01.1 | PE-CF594 |
| CD38 | OKT | PE |
